# Long-term single molecule localization microscopy uncovers dynamic co-assembly of Lrp6 and Ror2 into Wnt-signalosomes

**DOI:** 10.1101/2024.06.18.599024

**Authors:** Michael Philippi, Julia Dohle, Isabelle Watrinet, Michael Holtmannspötter, Jinye Li, Oliver Birkholz, Yi Miao, Ulrich Rothbauer, K. Christopher Garcia, Rainer Kurre, Jacob Piehler, Changjiang You

## Abstract

The conserved Wnt signaling has been classified as two categories of canonical and noncanonical Wnt signaling. With a high promiscuity of Wnt signaling, how receptors from the two distinct pathways re-arrange in multi-protein signalosomes remains elusive. We here developed single-molecule tracking and localization microscopy based on labeling with reversibly binding nanobodies (rbTALM) for imaging receptor dynamics in the plasma membrane for extended time periods. To this end, we engineered nanobody-tag pairs with fine-tuned binding stabilities ensuring single-molecule tracking with high fidelity, yet continuous exchange of photobleached labels. Multicolor rbTALM imaging enabled simultaneous tracking and super-resolution imaging of three different Wnt co-receptors in the same cell for more than one hour at video rate. Time-lapse correlation analyses uncovered cooperative association of canonical and noncanonical Wnt co-receptors into a common, hybrid Wnt signalosome, demonstrating the exciting possibilities of rbTALM imaging for exploring nanoscale dynamics across millisecond to hour timescales.

## Introduction

Wnt signaling plays a crucial role in stem cell differentiation and tissue organogenesis [1]. In humans, there are 19 different Wnt proteins, which are recognized by 10 different Frizzled receptors in conjunction with other co-receptors (PM) to activate Wnt signaling [2, 3]. Wnt signaling has been classified as β-catenin-dependent (canonical) and β-catenin-independent (noncanonical) Wnt signaling pathways [4] (**Fig. 1A)**. In the canonical Wnt signaling pathway, Wnts (e.g. Wnt3a) bind Frizzled (Fzd) to crosslink the co-receptor low-density lipoprotein receptor-related protein 5/6 (Lrp5/6) and trigger phosphorylation of the intracellular domain of Lrp5/6. This leads to recruitment of cytosolic scaffold proteins dishevelled (Dvl) and Axin, which are core components of the β-catenin destruction complex that furthermore includes the glycogen synthase kinase-3 (GSK3) and casein kinase 1 (CK1) [5]. Wnt-induced inhibition of the destruction complex leads to an increased level of the transcription factor β-catenin in the nucleus and the activation of gene expression [6].

**Fig. 1.**
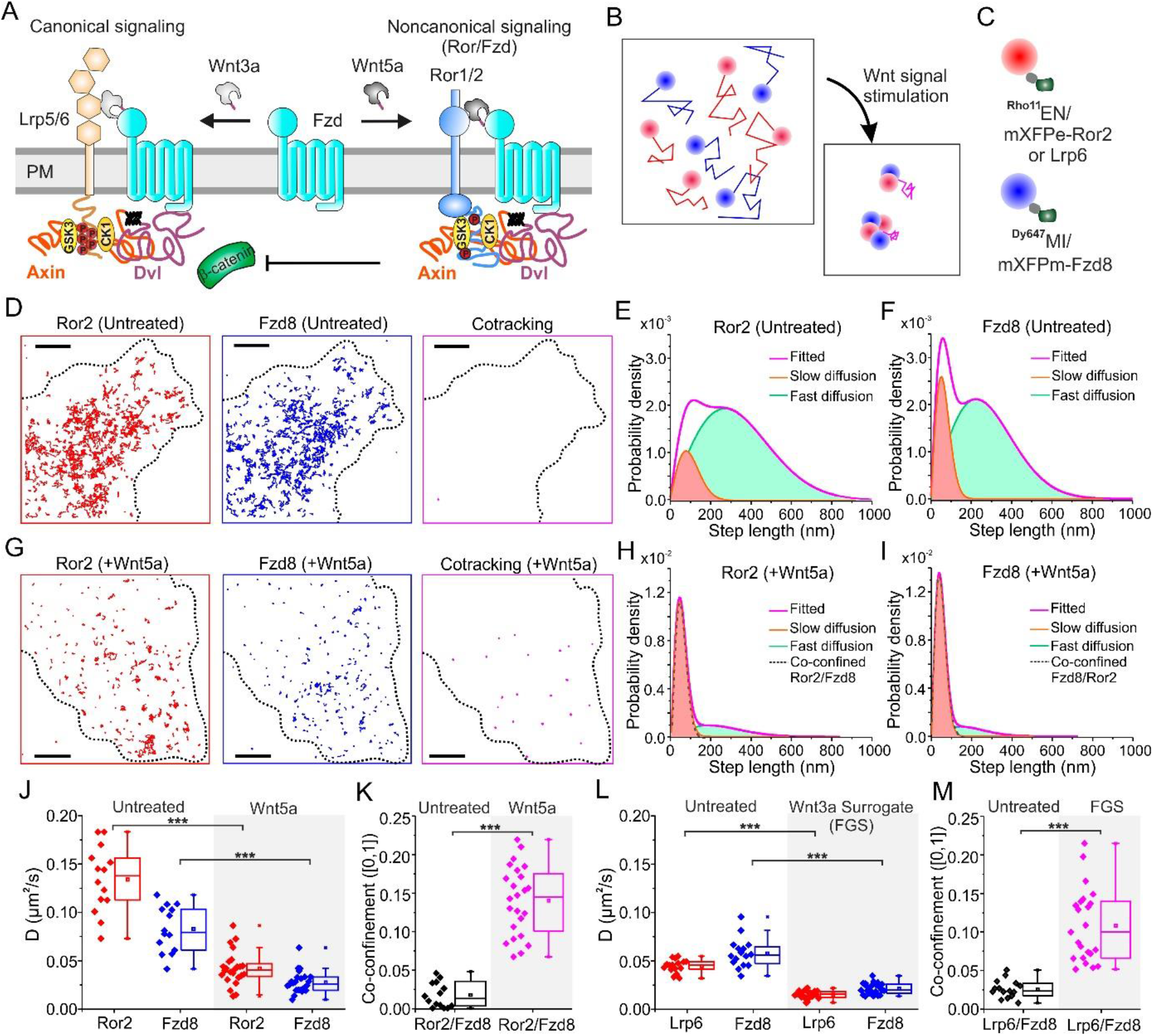
Receptor co-confinement as a hallmark of canonical and noncanonical Wnt signalosomes. (A) Scheme depicting receptors and effector interactions involved in the canonical and noncanonical Wnt signaling pathways. (B) Cartoon illustrating two-color single-molecule tracking of co-receptors upon Wnt signal stimulation. Binding and clustering of the receptors lead to co-confinement. (C) Scheme of stably labeling of mXFPe-Ror2/Lrp6 (red) and mXFPm-Fzd8 (blue) by ^Rho11^EN and ^Dy647^MI nanobodies, respectively. (D) Single-molecule trajectories of Ror2 (red) and Fzd8 (blue) of a representative cell before Wnt5a treatment with minimum receptor co-confinement (magenta). Scale bar: 5 µm. (E, F) Step length histogram indicating dominant fast diffusion fractions of Ror2 (E) and Fzd8 (F) at the untreated state. Number of steps in (E): 6450 and (F): 8845. (G) Trajectories of Ror2 (red) and Fzd8 (blue) as well as Ror2/Fzd8 co-confinement (magenta) of a representative cell after Wnt5a treatment. Scale bar: 5 µm. (H, I) Step length histogram showing decreased fast diffusion of Ror2 (H) and Fzd8 (I) and increase of slow diffusion after Wnt5a stimulation. Dash lines show the slow diffusion overlaps with normalized step length distribution of co-confined receptor. Number of steps in (H): 8874 and (I): 8606. (J) Average diffusion constants of Ror2 and Fzd8 before (*n =* 14 cells) and after Wnt5a stimulation (*n =* 24 cells). (K) Co-confinement of Ror2/Fzd8 after Wnt5a stimulation compared to untreated cells, i.e. (14.3 ± 4.7) % vs. (1.8 ± 1.7) %. Each dot represents the result of an individual cell. Significances of two-sample t-test: *** *p* < 0.001. (L) Average diffusion constants of Lrp6 and Fzd8 before (*n =* 16 cells) and after Wnt3a surrogate FGS stimulation (*n =* 24 cells). (M) Co-confinement of Lrp6/Fzd8 after stimulation vs. untreated, i.e., (10.7 ± 4.7) % vs. (2.7 ± 1.1) %. Significances of two-sample t-test: *** *p* < 0.001.

Tremendous efforts have been made to explore the molecular mechanism of canonical Wnt signaling [7-9]. The seven-pass Fzd transmembrane domain (Fzd-TMD) has a closed, small pocket in contrast to GPCRs [10, 11]. With a ∼40 aa flexible linker between Fzd-TMD and the Wnt-binding cysteine rich domain (Fzd-CRD), these imply that the canonical Wnt signaling is different to the ligand-induced allostery as in GPCR signaling. Activation by heterodimeric agonist Wnt surrogates [12, 13] or antibodies [14, 15] furthermore suggest that Wnt signaling follows a ligand-induced dimerization paradigm as represented by receptor tyrosine kinases (RTKs) activation [16]. In fact, four of 20 human RTK families are known to be co-receptors for noncanonical Wnt signaling [17, 18]. Receptor tyrosine kinase-like orphan receptor 1 and 2 (Ror1/2) are Wnt-binding RTKs involved in a noncanonical Wnt/Planar Cell Polarity (PCP) signaling axis [17, 19]. While originally known as ‘orphan’ receptor, Wnt5a was identified to be the ligand of Ror2 [20], and Ror1/2 have a Fzd-related CRD. However, structures of Ror2-CRD show it is either pre-occupied by a fatty acid [21] or lack the hydrophobic binding pocket [22]. This indicates Wnt5a needs to engage Fzds (e.g. Fzd2, 4, 5 or 8) [23, 24] to bind Ror2 and trigger its downstream signaling by GSK3 and the receptor kinase [25-27]. The structural insights support a competing binding model, in which different Wnt variants steer the Fzd-Dvl-Axin signaling axis to the canonical or the noncanonical pathway by either recruiting Lrp6 or Ror2 [24] (**Fig. 1A)**. Although the competing binding model well explains the Wnt5a antagonism of Wnt3a-induced canonical Wnt signaling, it is challenged by the surprising result that Wnt5a may also potentiate canonical Wnt signaling [28]. This apparent contradiction may be related to the additional complexity resulting from the assembly of “signalosomes” as the hallmark of Wnt signaling. The Wnt signalosome consists of multiple receptors and scaffold proteins [29-31]. Previous studies had highlighted the indispensable roles of the cytosolic scaffold proteins in formation of the canonical Wnt signalosomes [29, 32-34]. Recent studies of Axin1 and Dvl2 confirmed these intrinsically disordered scaffold proteins co-condensate into protein droplets [35-37]. Moreover, puncta of Ror2, Fzd and Dvl2 in the plasma membrane were observed in Wnt5a-stimulated noncanonical signaling [27, 38, 39]. These results collectively suggest that formation of Wnt signalosomes is a generic feature for both canonical and noncanonical Wnt signaling and co-condensation of cytosolic scaffold proteins plays a central role. The interplay between the different types of signalosomes at molecular level, however, has remained largely unclear.

To tackle this question, we have here developed a new approach enabling multi-color single-molecule localization microscopy in live cells to monitor spatiotemporal receptor dynamics during assembly of canonical and noncanonical Wnt signalosomes over extended time periods. For uncompromised single-molecule tracking with high temporal resolution during the course of signalosome assembly, we engineered tag-specific nanobodies with binding kinetics optimized for continuous exchange of fluorescence labels while maintaining labeling efficiency and complex stability. Thus, we achieved long-term super-resolved imaging of spatiotemporal receptor dynamics by tracking and localization microscopy (TALM) [40, 41]. We demonstrate that TALM imaging with reversible binders (rbTALM) can robustly quantify changes of receptor diffusion and nanoscopic receptor (co-)clustering without significant loss of labeling density over time. Thus, rbTALM imaging enabled tracking and super-resolution imaging of three Wnt co-receptors during sequential assembly of canonical and noncanonical signalosomes for more than 90 min at video rate. By applying spatial and spatiotemporal correlation analyses, we identified several key features governing Fzd8, Ror2 and Lrp6 co-clustering in a common Wnt signalosome.

## Results

### Characteristic co-confinement of Fzd8 and Ror2 in noncanonical Wnt signalosomes

We have previously explored signalosome assembly in the canonical Wnt signaling pathway by two-color single-molecule tracking of Lrp6 and Fzd8 [12, 13]. Using the same approach, we interrogated dimerization of Fzd8 and Ror2 as the key components of the noncanonical pathway (**Fig. 1B**). Leveraging our recently established orthogonal nanobody labeling technique [42], Ror2 and Fzd8 were fused with non-fluorescent variants of monomeric GFP, mXFPe and mXFPm, which are engineered for orthogonal labeling with the anti-GFP nanobodies “enhancer” (EN) and “minimizer” (MI), respectively (**Fig. 1C**). In the resting state, the diffusion of Ror2 and Fzd8 was dominated by the fast diffusion fraction with negligible heteromerization of the co-receptors as quantified by co-tracking analysis [12, 43] (**Fig. 1D-F**). Neither we could detect Ror2 homodimerization, which was probed by co-tracking of Ror2 labeled in two different colors (**Fig. S1A-D, Movie S1**). Both, the diffusion and intensity analyses showed Ror2 was monomeric (**Fig. S1E-G**) suggesting the absence of Ror2 pre-dimerization, which is in line with the previous findings of monomeric Fzd8 and Lrp6 at the resting state [12, 13]. Upon stimulation with Wnt5a, dramatically reduced mobility of Ror2 and Fzd8 was observed, with Ror2 and Fzd8 exhibiting strong co-confinement (**Fig. 1G-I, Movie S2**). Quantification from multiple cells revealed a decrease of the diffusion constants of Ror2 and Fzd8 by 60-70% (**Fig. 1J**) and 7-times increased co-confinement (**Fig. 1K**). The same tendency of strongly decreased diffusion constants and increased co-confinement was identified for nanobody-labeled Lrp6 and Fzd8 upon stimulation with the first-generation Wnt surrogate (FGS) (**Fig. 1L,M, Fig. S2, Movie S3**). This engineered cross-linker specifically dimerizes Fzd with Lrp6, and therefore selectively activates canonical Wnt signaling [12]. The observed puncta of Ror2/Fzd8 and Lrp6/Fzd8 upon stimulation with Wnt5a and FGS, respectively, highlighted formation of noncanonical and canonical Wnt signalosomes (**Movie S2, Movie S3**).

### Canonical and noncanonical Wnt signalosomes strongly overlap

To identify potential spatiotemporal correlation between the two types of signalosomes, we implemented three-color single-molecule tracking of Ror2, Lrp6 and Fzd8. To this end, HeLa cells were transiently transfected with Ror2, Lrp6 and Fzd8 *N*-terminally fused with mXFPe, HaloTag, and SNAP-tag, respectively, for orthogonal labeling with ^DY752^EN, ^TMR^HTL and SNAP Surface 647 (^647^SNAP) (**Fig. 2A**). In resting cells, each receptor showed typical free diffusion in the plasma membrane (**Fig. 2B**). Wnt stimulation significantly decreased receptor diffusion by adding either the mixture of Wnt3a & Wnt5a (**Fig. 2C**) or FGS & Wnt5a (**Fig. 2D**), which were quantified by analyses of diffusion constants (**Fig. 2E**) and co-confinement of Ror2/Fzd8, Fzd8/Lrp6 and Lrp6/Ror2 (**Fig. 2F**). Strikingly, more than one order-of-magnitude increased co-confinement of Lrp6/Ror2 (3.9% and 2.1% vs 0.2 %) was detected although the co-receptors are implicated in distinct Wnt signaling pathways (**Fig. 2F, Movie S4, Movie S5**). Overlapping of all three co-receptors could be clearly identified as co-clustered trajectories (**Fig. 2G-J, Fig. S3**). Taken together, these results suggest that co-stimulation of canonical and noncanonical Wnt signaling pathways results into co-clustering of Ror2, Lrp6 and Fzd8 into overlapping signalosomes. This finding thus challenges the receptor competing model, which predicts a mutual exclusion of Ror2 and Lrp6 for binding to the same co-receptor Fzd [24]. More detailed, time resolved analysis of co-receptor recruitment into Wnt signalosomes, however, was hindered by photobleaching, limiting single-molecule tracking to timespan of several seconds (**Fig. 2J**).

**Fig. 2.**
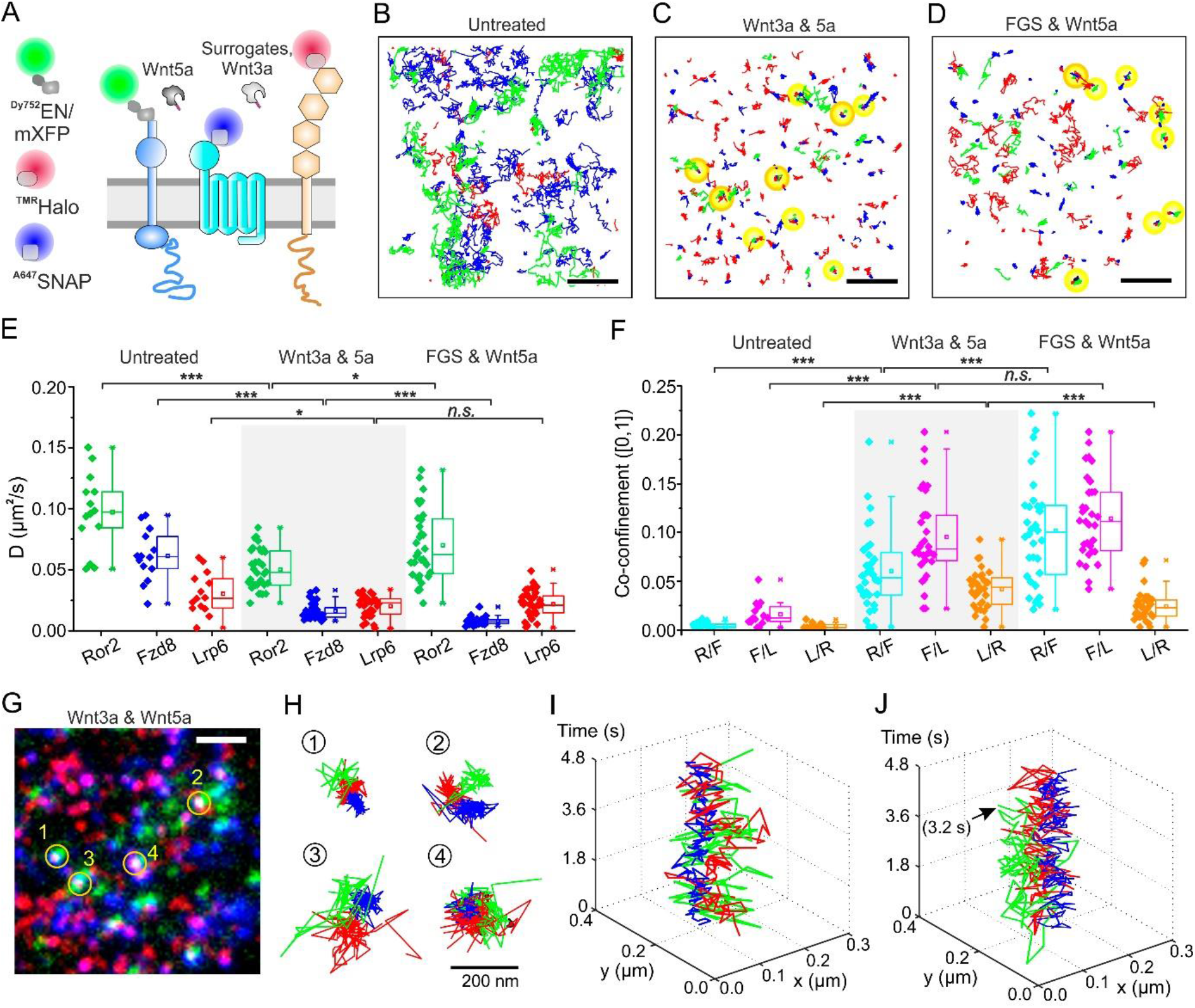
Rearrangement of Ror2, Lrp6 and Fzd8 by simultaneous canonical and noncanonical Wnt stimulation. (A) Scheme for orthogonal triple-color labeling of mXFP-Ror2 by ^Dy752^EN (green), HaloTag-Lrp6 by ^TMR^HTL (red) and SNAP-Fzd8 by ^A647^SNAP Surface (blue) for single-molecule tracking. Wnt3a and Wnt5a or FGS and Wnt5a were used for joint Wnt stimulation. (B, C, D) Representative trajectories of Ror2 (green), Lrp6 (red) and Fzd8 (blue) for untreated cells (B), in presence of Wnt3a & Wnt5a (C) and FGS & Wnt5a (D). Scale bars: 2 µm. Three-color co-confinement is marked in yellow. Trajectories in regions with orange outlines are shown in the following panels. (E) Average diffusion constants of Ror2 (green), Lrp6 (red) and Fzd8 (blue) in untreated cells (n = 14 cells), stimulated with Wnt3a & Wnt5a (gray bar, n = 32 cells), and stimulated by Wnt3a surrogate FGS & Wnt5a (n = 32 cells). (F) Co-tracking analyses of receptor pairs between Ror2 (*R*), Lrp6 (*L*) and Fzd8 (*F*) for the untreated state (*n* = 14 cells), stimulated with Wnt3a & Wnt5a (gray bar, *n* = 32 cells), and FGS & Wnt5a (*n* = 32 cells). Dots in the box chart represent result obtained from individual cells. Significances of two-sample t-test are: *** *p* < 0.001 and * *p* < 0.05. (G) Raw TIRF microscopy image of Ror2 (green), Lrp6 (red) and Fzd8 (blue) for single-molecule tracking analysis in (C). Regions containing colocalized Ror2/Lrp6/Fzd8 are marked with yellow circles. (H) Zoomed three-color co-localized trajectories. (I) Spatiotemporal coordinates of trajectories in the 4^th^ group. (J) Spatiotemporal coordinates of co-diffusion trajectories marked with orange outline in panel (D). Arrow shows the observed photobleaching step.

### Reversible nanobody enables long-term tracking and localization microscopy

To circumvent the limitation by photobleaching in long-term single-molecule tracking, we developed reversible binders for labeling to replenish bleached fluorophores (**Fig. S4A**). For this purpose, we engineered nanobody/target pairs having an intermediate dissociation rate constant (*k_d_* ∼0.1 s^-1^) and fast association rate constant (*k_a_* > 10^5^ M^-1^s^-1^). Thus, exchange rates of fluorophore matching the bleaching kinetics were achieved while maintaining sub-micromolar affinities to ensure good labeling efficiencies. For orthogonal labeling of proteins-of-interest, five nanobodies were screened (**Fig. S4B-G**) including (i) a rationally designed double-mutant ALFA nanobody M63A/R65E (ALFAnbAE) for reversible binding of the ALFAtag; (ii) an anti-HaloTag nanobody (HD10) directly selected from a nanobody library; (iii) an anti-maltose binding protein (MBP) “sybody” (MSyb) [44]; (iv) the anti-GFP nanobody EN that binds mCFP-H164N (CFP*HN*) with low affinity [45]; and (v) the anti-GFP nanobody MI for the engineered low-affinity target mXFPe [42]. After characterizing the binding affinities and kinetic constants in *vitro* (**Fig. S4G**), we examined the effective labeling at cell surface using engineered chimeric constructs containing SNAP tag and nanobody target as the model transmembrane receptors (**Fig. S4H**). The degree-of-labeling (*DOL*) of nanobodies were quantified by comparing to the well-characterized *DOL* of 43% for SNAP tag [46] in two-color single-molecule tracking (**Fig. S4I, J**). For all nanobodies tested, constant single-molecule localization counts were achieved during 4.2 min TIRF imaging at video rate (**Fig. S4K**), confirming efficient exchange of bleached fluorophores. Most robust labeling results were obtained with the binding pairs ALFAnbAE/ALFA-tag, HD10/HaloTag and EN/CFP*HN*, which therefore were selected for orthogonal labeling of the three Wnt co-receptors.

### Long-term rbTALM imaging uncovers nanoscale, condensate-like Wnt signalosomes

Leveraging reversible nanobody labeling, we carried out long-term single-molecule tracking and super-resolution imaging of canonical Wnt signalosomes by rbTALM imaging. For proof-of-concept experiments, Wnt signalosome formation in a HeLa cell line stably co-expressing HaloTag-Lrp6 and SNAP-Fzd8 was explored. The HaloTag was reversibly labeled by nanobody HD10 conjugated with Dy647 dye (^Dy647^HD10), which was also covalently labeled with TMR-HaloTag Ligand (^TMR^HTL) for comparison (**Fig. 3A**). Using the next generation surrogate (NGS) for stimulation, clustering of HaloTag-Lrp6 as the hallmark of canonical Wnt signalosome was visualized by TALM (**Fig. 3B-C**). NGS is a more recently developed cross-linker of Fzd and Lrp6, which is based on a DARPin that recognizes the CRD of Fzd competitive to Wnts [13]. In TIRF microscopy, specific binding of ^Dy647^HD10 to TMR-stained cells was observed with a density ideal for single-molecule tracking (**Fig. 3C, Fig. S5A-D**). While fluorescence intensity of the ^TMR^HTL channel continuously decreased due to photobleaching, the ^Dy647^HD10 channel showed no loss of localization counts, confirming efficient replenishing of bleached fluorophores (**Fig. 3D, Fig. S5E-F**). TALM super-resolution images were generated by rendering localized Lrp6 molecules accumulated over multiple frames using Gaussian blurring with the standard deviation equal to the average localization precision (**Fig. 3E-G**) [47]. Different ways of data processing yield either time-lapse or accumulated TALM images (**Fig. 3H**). Results from a typical TALM image within the region-of-interest (ROI) highlighted in **Fig. 3C** are shown in **Fig. 3I, J**. The rendered TALM image identifies several Wnt signalosomes (**Fig. 3I**), where the diffusion of Lrp6 is strongly confined (**Fig. 3J**, **Fig. S5G-H**). Within these signalosomes, however, Lrp6 clearly showed mobility (**Fig. S5I-L**, **Movie S6**), in line with condensate-like properties of these clusters, which we and others have recently proposed [37, 48, 49]. We further quantified Lrp6 confinement by pair correlation analysis [43, 47] (**Fig. 3K**) that yielded a relatively high confinement probability (*cp*) of 0.68 and confined radius (*λ*) of 160 nm (**Fig. 3L**), in line with the strong Wnt signalosome formation upon NGS stimulation [13].

**Fig. 3.**
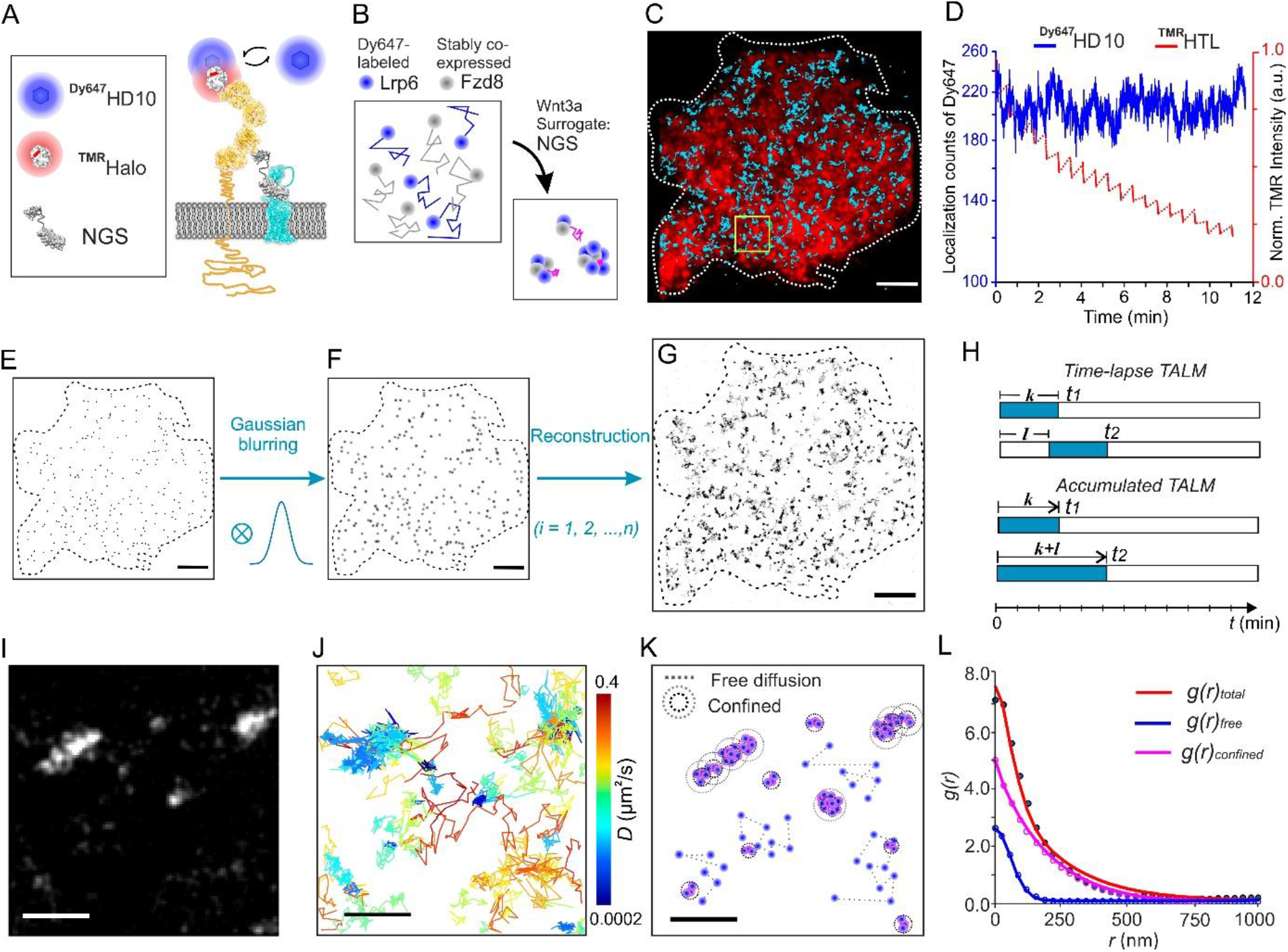
Reversible nanobody labeling of Wnt receptors for long-term rbTALM. (A) Proof-of-concept of reversible nanobody binding of HaloTag-fused Lrp6 by ^Dy647^HD10 (blue) in comparison to covalent ligand labeling by TMR (red). Agonist Wnt surrogate NGS is used for stimulation. Structures are adapted from Protein Data Bank (PDB): HaloTag (PDB: 5Y2Y), NGS (PDB: 6NDZ), Lrp6 (PDB: 3S94 and 3S8Z), and Fzd (PDB: 4F0A and 6WW2). (B) Cartoon illustrating changes in diffusion and confinement of ^Dy647^HD10-labeled HaloTag-Lrp6 as the indication of canonical Wnt signalosome formation. (C) Merged TIRF microscopy image of the TMR channel (red) with single-molecule trajectories of ^Dy647^HD10-labeled HaloTag-Lrp6 (cyan) in a living cell. Dash line marks cell boundary. Scale bar: 5 µm. (D) Single-molecule localizations per frame of ^Dy647^HD10 (blue) and mean fluorescence intensity of TMR channel (red) versus time. ^Dy647^HD10 shows an average density of 0.3 molecules/µm^2^ in the cell. Up-and-down of the intensity in TMR channel is due to fluorescence recover by the pause of excitation. (E-H) Illustration of image processing for TALM reconstruction. (E) Localizations of ^Dy647^HD10-labeled HaloTag-Lrp6 (black dots) obtained from a single image frame. Dashed line indicates cell. (F) Each localization is blurred by convolution with mean localization precision (Gaussian). (G) Accumulated frame-by-frame TALM reconstruction of Lrp6 based on 300 frames. Scale bars (E-G): 5 µm. (H) Time-lapse TALM versus accumulated TALM reconstruction: In time-lapse TALM, a sliding observation window (𝑘 frames, turquoise bar) as well as a step size (𝑙 frames, blank bar) is defined to bin localizations with a specific time interval (*t*_1_, *t*_2_, etc.). These localizations can be accumulated to obtain accumulated TALM stacks with the same temporal resolution. (I) Accumulated TALM image of the region-of-interest (ROI) highlighted in panel C reconstituted from 8,460 Lrp6 localizations obtained in 84 s. (J) Single-molecule trajectories of Lrp6 obtained in 84 s in the same ROI. Color-coded by diffusion constant (logarithmic scale). (K) Schematic illustration of pair correlation analysis for TALM image in panel (I). Confined diffusion leads to convolution of localization PSF as clusters (dash circles). Scale bars (I-K): 500 nm. (L) Confinement probability (*cp*) of Lrp6 quantified by pair correlation analysis.

### Kinetics of nanoscale Wnt signalosome assembly captured at video-rate time resolution

To unambiguously resolve the process of Wnt signalosome assembly, we conducted two-*color* rbTALM imaging of HeLa cells, which were co-expressing HaloTag-Lrp6 and CFP*HN*-Fzd8 and orthogonally labeled with ^Rho11^HD10 and ^Dy647^EN (**Fig. 4A**). TIRF microscopy of the same cell was carried out continuously for 10 min before and 60 min after adding NGS (120,000 frames at video rate acquisition). Quantification of receptor diffusion and co-confinement in the same cell before adding NGS and at the very end of the experiments (**Fig. 4B-C, Fig. S6A-F**) yielded comparable results as obtained in the multi-cell assays shown in **Fig. 1L, M**. The constant localization counts of Lrp6 and Fzd8 over 1 h confirmed successful two-color rbTALM imaging (**Fig. 4D**). By selectively rendering co-localized Lrp6/Fzd8 throughout the 70 min experiment, we generated an overview colocalization TALM (coloc-TALM) image highlighting Wnt signalosome formation in response of NGS-stimulation (**Fig. 4E**). By reconstructing coloc-TALM images of Lrp6 and Fzd8 in the time-lapse mode with a step size of 1.75 min, the process of Lrp6/Fzd8 assembling into signalosomes can be followed at the level of individual receptor pairs (**Fig. 4F, H, Fig. S6G, H**). Under these conditions, disappearing of clusters can be ascribed to transient, substoichiometric labeling of receptors used in these experiments. By contrast, the accumulated coloc-TALM images captures the entire process of signalosome assembly (**Fig. 4G, I, Fig. S6I, J**). Strikingly, the accumulated coloc-TALM images revealed formation of small seed cluster in the early phase which over time grow into larger oligomers (**Fig. 4I, Fig. S6J**). These observations of growth and merging of receptor clusters are in line with nanoscale protein condensation being responsible for signalosome assembly (**Movie S7**). Applying pair correlation analysis to the time-lapse coloc-TALM images furthermore enabled quantifying the kinetics of signalosome assembly (**Fig. S7A-F**). Mono-exponential fitting of co-confinement probabilities (*ccp*) *vs* time resulted to a time constant of 20 ± 12 min (**Fig. 4J**). The unchanged basal confinement radius (λ) of 160 ± 60 nm can be ascribed to the diffusional confinement of individual receptors, in line with the ∼300 nm-diameter native compartmentalization in the plasma membrane of HeLa cells we have previously observed by TALM of cytokine receptors [43] (**Fig. 4K**). These results confirmed residual receptor mobility within signalosomes in line with a process driven by protein condensation. Pair correlation analysis of the accumulated coloc-TALM images resulted to the similar time constant of 22.1 ± 2.9 min for signalosome assembly with gradually increased cluster sizes upon NGS stimulation (**Fig. 4L, M, Fig. S7G, H**). After 30 min stimulation, the λ values reached a level of 302 ± 24 nm and remained constant until the final stage, indicating an average signalosome diameter of 600 nm. However, accumulated TALM images only reflect the growth of signalosome, but cannot account for loss or re-arrangement. To avoid this bias, we extracted the kinetics of signalosome formation from pair correlation analysis of time-lapse TALM images, which do not sample the entire signalosome, but much more reliably report overall changes.

**Fig. 4.**
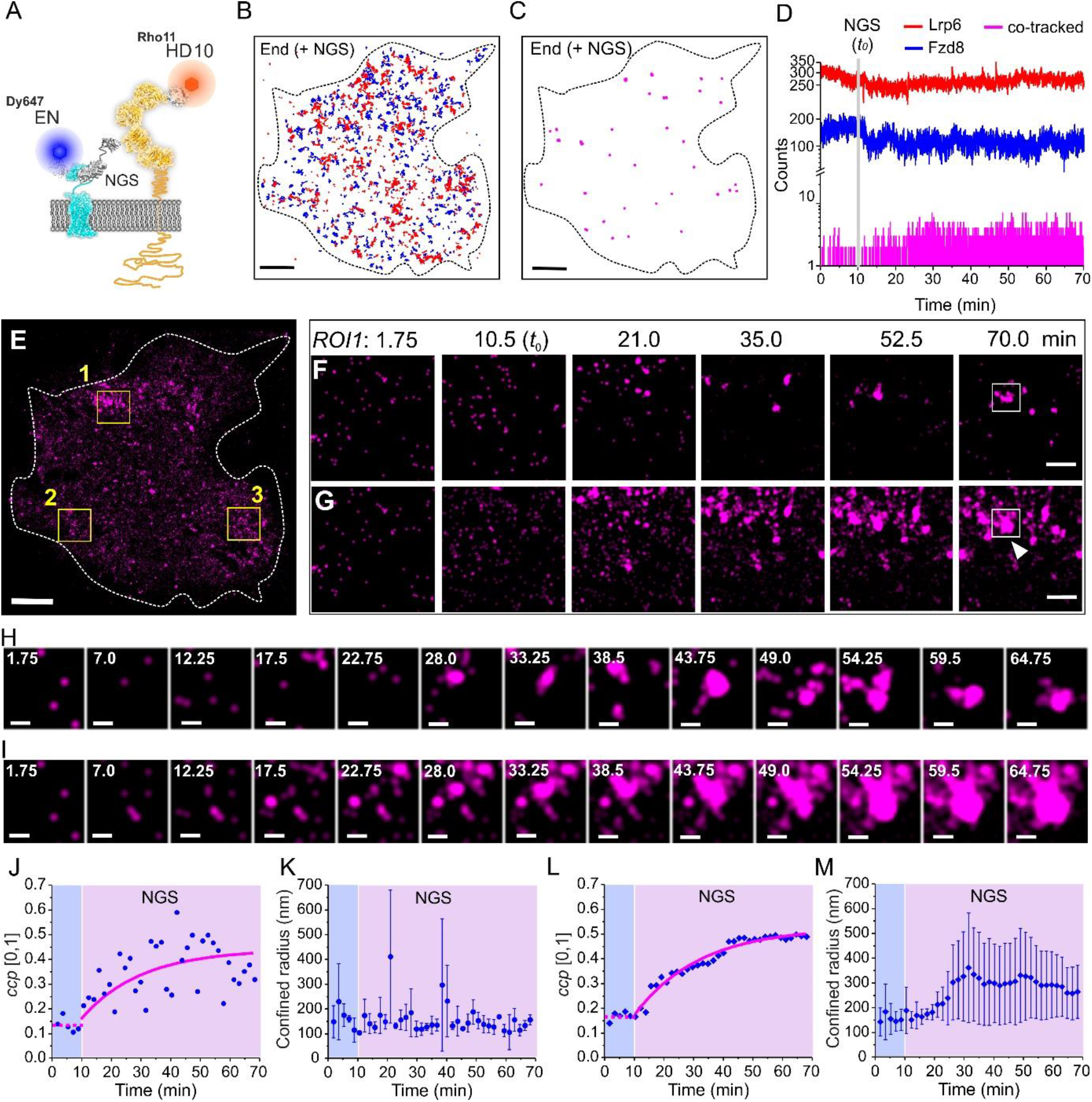
Formation of canonical Wnt signalosome monitored by two-color co-localization TALM (coloc-TALM). (A) Orthogonal and reversible nanobody labeling of HaloTag-Lrp6 and CFP*HN*-Fzd8 with ^Rho11^HD10 (red) and ^Dy647^EN (blue), respectively. Agonist Wnt surrogate NGS used for stimulation. (B) Live cells were treated with NGS and imaged over 70 min at video rate (120,000 frames). Trajectories of Lrp6 (red) and Fzd8 (blue) at the end of the experiment based on 150 frames. Dashed line indicates cell shape. (C) Corresponding co-tracking trajectories (>10 frames) of Lrp6/Fzd8. Scale bars: 5 µm. (D) Localization per frame versus time of Lrp6 (red), Fzd8 (blue) and co-tracked Lrp6/Fzd8 (magenta). Gray line marks the starting time (*t*_0_) of stimulation with NGS. (E) Accumulated coloc-TALM image showing Lrp6/Fzd8 co-confinement zones (magenta) based on 12,000 frames with an interval of 350 ms. Scale bar: 5 µm. (F, G) Time-lapse TALM images (F) and accumulated TALM images (G) of ROI1 in panel (E). Scale bars: 1 µm. Image series show co-localized Lrp6/Fzd8 (magenta) in a 500 frames observation window (2.92 min) sliding at a step size of 300 frames (1.75 min). Selected time points including 𝑡_0_ are indicated. (H) Temporal Lrp6/Fzd8 co-clustering highlighted for the region indicated by the white rectangle in panel (F), and (I) indicated by white rectangle/triangle in panel (G). Scale bars: 200 nm. Numbers are the step time in min. (J) Pair correlation analysis of time-lapse coloc-TALM for quantifying kinetics of Lrp6/Fzd8 co-clustering upon NGS stimulation. Co-confinement probabilities (*ccp*) are mean values from the three ROIs indicated in panel (E). Magenta curve indicates mono-exponential fit. Dash magenta line shows the mean values before stimulation. Untreated state (aqua blue) and NGS stimulation (pink) are marked with color bars. (K) Confinement radius of three ROIs plotted as mean ± sd versus time. (L) Pair correlation analysis of accumulated TALM images. *ccp* are mean values obtained from three ROIs. Magenta curve indicates mono-exponential fit. (M) Confinement radius of three ROIs plotted as mean ± sd versus time.

### Wnt5a triggers Ror2 co-recruitment into common Wnt signalosomes

To decipher the interplay of canonical and noncanonical Wnt signalosome assembly, we applied three-color rbTALM imaging to simultaneously capture the co-organization and spatiotemporal dynamics of Fzd8, Lrp6 and Ror2. To this end, ALFA-Ror2, HaloTag-Lrp6 and mCFP*HN*-Fzd8 were co-expressed in HeLa cells for orthogonally reversible labeling with ^Rho11^AnbAE, ^Dy647^HD10 and ^Dy752^EN. After sequentially adding NGS and Wnt5a, three-color single-molecule co-tracking confirmed significantly increased pairwise co-confinement of Ror2, Lrp6 and Fzd8 (**Fig. 5A-D, Fig. S8A-D, Movie S8**). Having confirmed the viability of the system, we monitored the entire process of sequential assembly of canonical and noncanonical signalosomes by continuous three-color TIRF imaging of a single cell for 94 min at video rate image acquisition. Frame-by-frame single-molecule localization confirmed that labeling density in all three channels was robustly maintained throughout the entire experiment (**Fig. S8E, F, Movie S9**). The accumulated TALM image identifies co-clustering of all three receptors (**Fig. 5E, Fig. S8G-I**), confirming overlapping of canonical and noncanonical Wnt signalosomes. Interestingly, co-clustering was most pronounced at the cell periphery (**Fig. S8A, E**), which may suggest a contributing role of the cortical actin cytoskeleton, but could also be caused by effective photobleaching being more pronounced underneath the cell due to diffusion-limited rebinding rates. We therefore selected ROIs at the cell rim to analyze the spatiotemporal re-arrangement of the Wnt co-receptors using the established protocol [47, 50] (**Methods**). In these regions, decrease of co-receptor mobility in line with the stimulating agent was observed, yet with different kinetics (**Fig. 5F, Fig. S9A-F**). A rapid decrease of the diffusion constants of Lrp6 and Fzd8 with a time constant of ∼2 min was observed after addition of NGS, followed by a slow further decline during the following 30 min down to < 20% of the original level (**Fig. 5F**). Interestingly, the diffusion constant of Ror2 also decreased directly after NGS stimulation, yet only mildly (by ∼20%). Strikingly, transient confinement of Ror2 in NGS-induced signalosomes was observed (**Fig. 5G, H, Movie S10**), which can explain this effect and confirmed interaction of Ror2 with canonical signalosomes. After adding Wnt5a, a further, very slow decrease of the Ror2 diffusion constant to < 50% of the original level was observed (**Fig. 5F**).

**Fig. 5.**
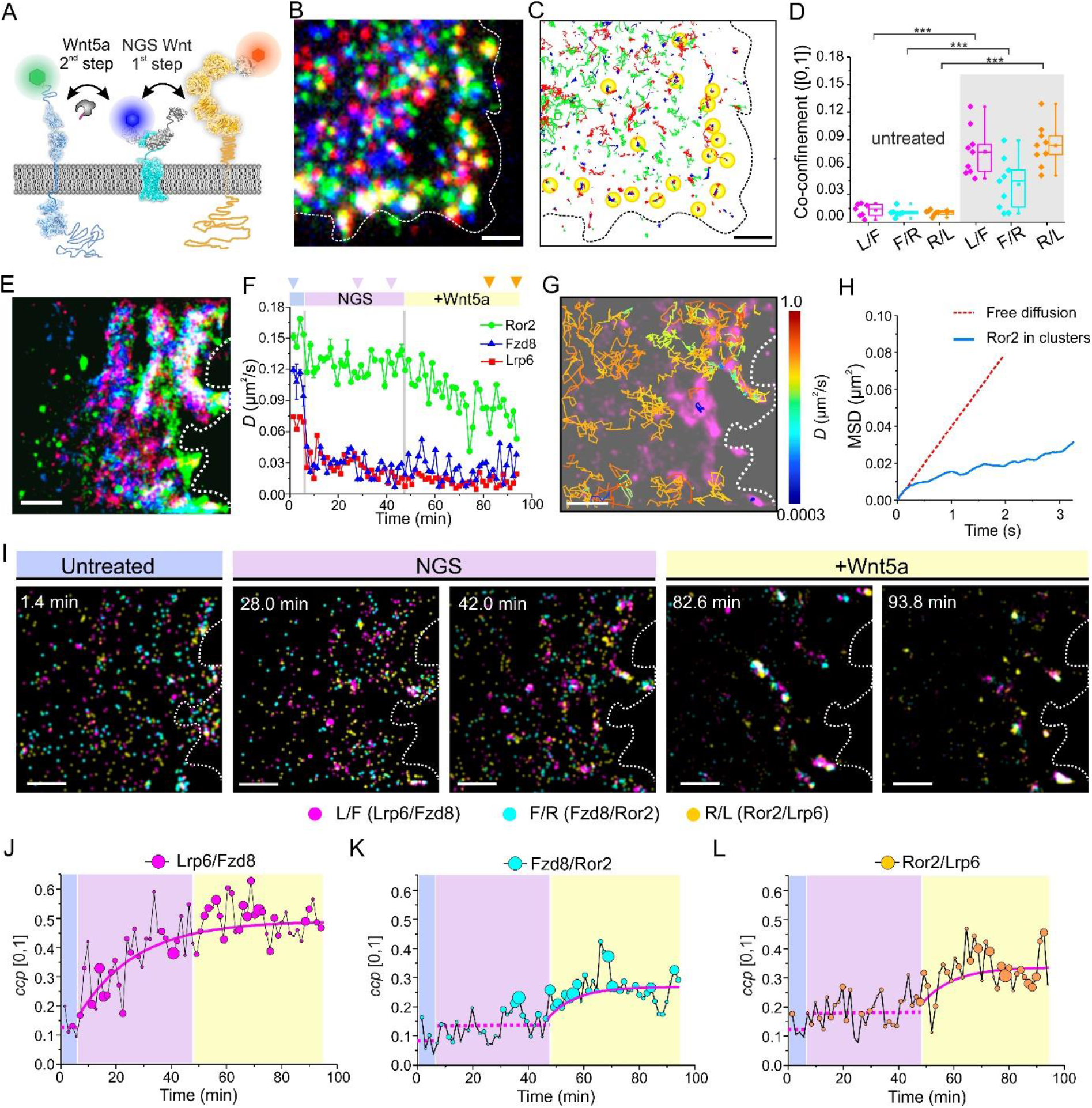
Spatiotemporal reorganization of co-receptors during canonical and noncanonical Wnt signaling. (A) Scheme of sequential activation for canonical and noncanonical Wnt signaling by NGS and Wnt5a. ALFA-Ror2, HaloTag-Lrp6 and CFP*HN*-Fzd8 were co-expressed in HeLa cells and reversibly labeled with ^Rho11^AnbAE, ^Dy647^HD10 and ^Dy752^EN, respectively. Structure of Ror2 is adapted from PDB (7ME4, 6OSV and 3ZZW). (B-C) Long-term triple-color TIRF imaging (160,840 frames, 94 min) at video rate of Ror2 (green), Lrp6 (red) and Fzd8 (blue) during sequential stimulation by NGS and Wnt5a. (B) Image shows crop of merged TIRF microscopy channels at end of experiment (cf. full field of view in Fig. S8A). Dashed line indicates cell shape. (C) Single-molecule trajectories of all three receptors shown in panel (B) in the last 300 frames. Yellow cycles mark co-confinement zones. Scale bars (B, C): 2 µm. (D) Co-tracking analyses of receptor pairs between Ror2 (*R*), Lrp6 (*L*) and Fzd8 (*F*) before (n =7) and after sequential Wnt stimulation (n = 9 cells). *** p<0.001 for two-sample t-test. (E) Accumulated three-color rbTALM of a representative ROI at the cell boundary (cf. Fig. S8E ROI2) showing Ror2 (green), Lrp6 (red) and Fzd8 (blue). TALM image is based on 20,105 frames with an interval of 280 ms. Scale bar: 1 µm. (F) Change of diffusion constants versus time for each receptor in the area shown in (E). Gray lines indicate the time of adding NGS (5.8 min) and Wnt5a (47.1 min). Arrows indicates the acquisition time of images shown in panel (I). (G) Trajectories of Ror2 after adding NGS but before Wnt5a (46.9 min) overlaid onto the corresponding Lrp6/Fzd8 coloc-TALM image (magenta). Color bar indicates diffusion constants of shown trajectories (logarithmic scale). (H) Mean squared displacement (MSD) of pooled Ror2 trajectories (150 frames) overlapped within Lrp6/Fzd8 clusters. MSD of free diffusion (red dash line) is shown for comparison. (I) Time-lapse three-color co-localization TALM images of different receptor pairs at different stages of stimulation. Scale bars (E-I): 1 µm. Co-localized Lrp6/Fzd8 (magenta), Fzd8/Ror2 (cyan), and Ror2/Lrp6 (yellow) are shown. Untreated state (aqua blue), NGS stimulation (pink) and NGS+Wnt5a stimulation (yellow) are marked with color bars, respectively. (J-L) Co-confinement probabilities (*ccp*) versus time for Lrp6/Fzd8 (J), Fzd8/Ror2 (K) and Ror2/Lrp6 (L) at different status. Bubble plots are based on mean values of the three ROIs marked in Fig. S8E. Size of plots represents standard deviation. Mono-exponential fits (solid magenta lines) and overall mean values (dashed magenta lines) are shown.

### Canonical signalosomes alter the pathway of noncanonical signalosome assembly

Accumulated TALM images confirmed strong co-clustering of all three co-receptors during the entire course of the experiment. The spatiotemporal dynamics of signalosome assembly was further analyzed by three-color time-lapse TALM of the co-localized receptors (**Fig. 5I, Fig. S9G-I, Movie S11**) and pair correlation analysis (**Fig. S9J-L**). The characteristics of canonical signalosome assembly upon NGS-stimulation nicely matched the results presented above, with a Fzd8/Lrp6 co-clustering time constant of 20.6 ± 5.4 min (**Fig. 5J, Fig. S9J**). After addition of Wnt5a, significant overlap of co-clustered Lrp6/Fzd8, Fzd8/Ror2 and Ror2/Lrp6 were observed in the three-color colocalization TALM image (**Fig. S9G-I**). To discriminate possible interactions between receptors, we deconvolved the three-color coloc-TALM images into three pairs of two-color coloc-TALM (**Fig. S10)**. Interestingly, the two-color time-lapse coloc-TALM images clearly uncovered Wnt5a-induced partitioning of Ror2 into canonical signalosomes, while co-clustering of Lrp6/Fzd8 hardly changed (**Movie S12, S13**). Indeed, very similar time constants were observed for co-clustering of Ror2/Fzd8 (𝜏: 11.4 ± 4.8 min) and Ror2/Lrp6 (𝜏 : 10.2 ± 4.6 min) (**Fig. 5K, L, Fig. S9K, L**). In control experiments, the changes in the spatiotemporal dynamics of Lrp6, Fzd8 and Ror2 upon direct stimulation for noncanonical Wnt signalosome assembly was explored by three-color rbTALM imaging (**Fig. 6A, Fig. S11A-C**). Selective clustering of Fzd8/Ror2 was observed (**Fig. 6B**), accompanied by strongly decreased mobility of Ror2 (∼80 %) and Fzd8 (∼50 %), while Lrp6 showed a mildly decreased diffusion constant (25 %) (**Fig. 6C, Fig. S11D-G**). However, much faster decay of Ror2 mobility was observed as compared Wnt5a stimulation after formation of canonical signalosomes. Pair correlation analysis of the time-lapse coloc-TALM images confirmed a Wnt5a-triggered time-dependent increase of Fzd8/Ror2 co-confinement with a time constant of 23.8 min (**Fig. 6D, Fig. S11H-N**). By contrast, negligible changes in Ror2/Lrp6 or Lrp6/Fzd8 co-clustering was observed (**Fig. S11O, P**).

**Fig. 6.**
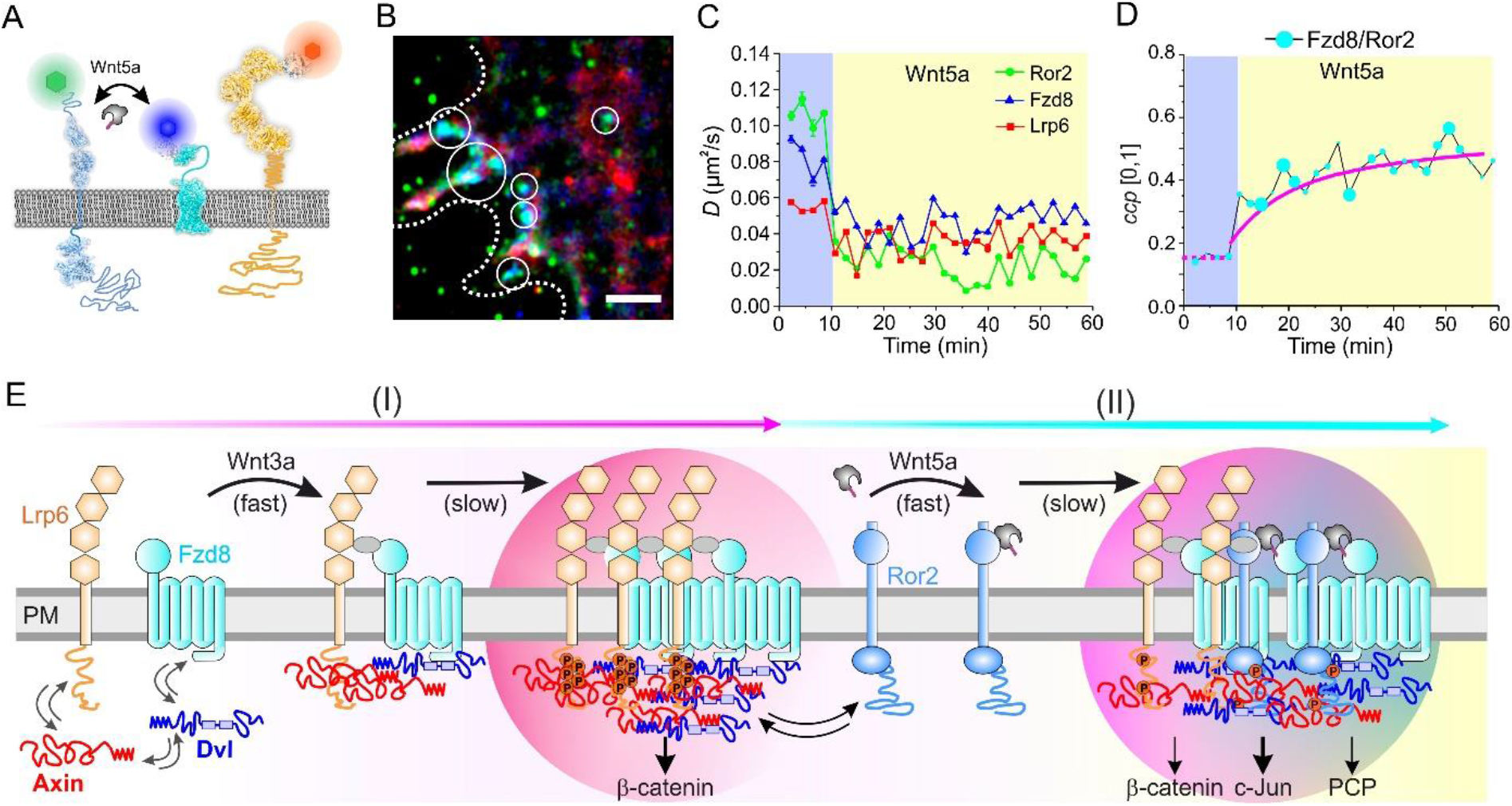
Noncanonical Wnt signalosome and the common Wnt signalosome. (A) Scheme of inducing noncanonical Wnt signaling by Wnt5a in cells co-expressing canonical and noncanonical Wnt receptors. (B-D) Long-term triple-color TIRF imaging (100,000 frames, 58 min) at video rate of Ror2 (green), Lrp6 (red) and Fzd8 (blue) during stimulation Wnt5a (cp. Fig. S11). (B) Representative cropped TALM image of Ror2 (green), Fzd8 (blue) and Lrp6 (red) after Wnt5a stimulation (ROI3 in Fig. S11B). TALM image was reconstituted based on 12,500 frames with interval of 280 ms. White circles mark the clustering of Fzd8 and Ror2 as the hallmark of noncanonical Wnt signalosome. Dashed line shows cell boundary. Scale bar: 1 µm. (C) Diffusion constants versus time for each receptor. (D) Time-lapse pair correlation analysis of receptor co-clustering. Co-confinement probability (*ccp*) of Fzd8/Ror2 are plotted as mean values of three ROIs (see Fig. S11). Size of plots represents standard deviation. Kinetics was fitted by mono-exponential function (magenta line). (E) Schematic model illustrating stepwise formation of the canonical Wnt signalosome and its transition into the common Wnt signalosome. In step (I), freely diffusing Lrp6 and Fzd8 rapidly form dimers upon NGS stimulation. This triggers the formation of canonical Wnt signalosome via co-condensation of Axin1 and Dvl2 (black dash arrow) to activate transcription regulation via β-catenin. Ror2 is already transiently trapped (blue dash arrow) in the canonical signalosome due to weak interaction with Axin/Dvl co-condensates, facilitating transition into the common signalosome upon Wnt5a binding (step II). Transition kinetics is probably limited by the stability of the Fzd8, leading to the activation of additional pathways.

## Conclusions

Live-cell single-molecule imaging offers unique insights to resolve the spatiotemporal organization of biomolecules at nanoscale. Most strikingly, individual biomolecules can be tracked within their biological context, providing important information on assembly and dynamics of supramolecular complexes and underlying cellular structures. Such approaches require fast image acquisition at relatively high excitation power densities, causing relatively fast photobleaching. Therefore, overall acquisition times of single-molecule tracking experiments are typically limited to several seconds, obstructing application to slow cellular processes. sptPALM and related photoswitching-based techniques [51] are useful to sequentially probe spatiotemporal dynamics of proteins expressed at high densities. Long-term single-molecule tracking at more physiological expression levels over the course of minutes to hours, however, has so far not been achieved. To overcome this limitation, we have here introduced single-molecule tracking and localization microscopy with reversible binders (rbTALM) to replenish bleached labels during image acquisition. We found that binding affinities of *K_D_* = 100 – 500 nM and complex lifetimes of ∼10s were most suitable for this approach, ensuring good labeling efficiencies, high tracking fidelities and exchange kinetics similar to typical photobleaching rates. Strikingly, we achieved video-rate rbTALM imaging in living cells at up to three channels with no significant loss in labeling density and uncompromised tracking fidelity during acquisition at video rate over up to 90 min. The rbTALM imaging technique is conceptionally related to live-cell PAINT approaches based on transient labeling of peptide/protein-subdomain [52-55] or DNA/RNA [56-58] binding pairs. However, the binders used for rbTALM were engineered for longer lifetimes and higher affinity to achieve not only extended single-molecule trajectories, but also higher degree of labeling required for effective co-localization analyses. We here already demonstrate the capability for long-term rbTALM imaging with three different colors, but including a fourth channel will be straight forward [42]. These capabilities make rbTALM imaging a unique approach for exploring the spatiotemporal dynamics of Wnt signalosome assembly, which involves fast (second-scale) and slow (hour-scale) processes.

Leveraging the capability of multi-color rbTALM imaging, we here focused on the interplay of canonical and noncanonical Wnt signaling by monitoring receptor diffusion and co-organization over more than an hour. Our observations uncovered several intriguing features of stepwise formation of a common Wnt signalosome, as summarized in the scheme of **Fig. 6E**. Stimulation of canonical signalosome formation by NGS induced rapid dimerization of freely diffusing Lrp6 and Fzd8, leading to dramatic reduction in mobility within 2 min. By contrast, co-clustering of Lrp6/Fzd8 into distinct signalosomes took place one order-of-magnitude slower timescale (∼20 min), exhibiting distinct features of a nucleation-growth process. These characteristics are well in line with recent models suggesting receptor dimerization triggering the formation of canonical Wnt signalosomes by co-condensation with the scaffold proteins Axin and Dvl [35, 37]. Strikingly, Ror2, the co-receptor for noncanonical Wnt signaling, which is not recognized by NGS, was found transiently trapped in NGS-induced canonical signalosomes. This can be explained by weak interactions of Ror2 with the Axin/Dvl co-condensates in the canonical signalosomes. Subsequent stimulation with Wnt5a promoted strong co-clustering of Ror2 with Fzd8 and Lrp6 in a common Wnt signalosome. Wnt5a-induced partitioning of Ror2 into common signalosomes was much slower as compared to dimerization of Ror2 and Fzd8 upon direct stimulation of noncanonical Wnt signaling. We ascribe this to Wnt5a and NGS competing for an overlapping binding pocket of Fzd8 [13] (**Fig. S12**), excluding NGS and Wnt5a binding to the same Fzd8. Thus, formation of the common signalosome was not caused by Wnt-mediated cross-linking of the co-receptors, but to Axin-Dvl co-condensates attracting Ror2 and promoting local recruitment of Fzd8. The kinetics of this process is limited by the stability of the NGS-Fzd8 complex [59], which probably is higher as compared to the corresponding Wnt3a-Fzd8 complex. With the central role of Axin-Dvl co-condensates in spatially defining signalosome assembly, simultaneous co-stimulation of canonical and noncanonical Wnt signaling yielded very similar common signalosomes. Taken together, our results support the emerging concept of a common canonical and noncanonical signalosome that is fundamentally driven by co-condensation of the cytosolic scaffold proteins Axin and Dvl. Rather than the original dichotomic concept, the formation of a common signalosome supports an integrated, multi-tiered perspective of canonical and noncanonical Wnt signaling [60, 61]. Further studies focusing on the composition and dynamics of the cytosolic signaling complex will be required to shed light into the underlying regulatory mechanisms. Extending rbTALM imaging to intracellular components of the Wnt signalosome, e.g. by exploiting reversible HaloTag labeling [62, 63], can be envisioned to provide an in-depth understanding about the cooperativity of receptor assembly and cytosolic protein condensation during Wnt signaling.

## Acknowledgements

We thank Markus Seeger in University of Zürich for providing the plasmid of anti-maltose binding protein sybody. We thank H. Kenneweg, A. Budke-Gieseking, G. Hikade and W. Kohl for the excellent technical assistance. This project is supported by funding to C.Y., J.P., and R.K. from the DFG (YO 166/1-1, SFB 1557, projects 13 and Z2, and by RTG 2900). K.C.G. is an investigator with the Howard Hughes Medical Institute and is supported by the Ludwig Foundation.

## Author contributions

J.P. and C.Y. conceived the project. M.P. performed the rbTALM of Wnt signaling and analyzed the results. J.D. conducted single-molecule tracking of Wnt receptors and characterized nanobody binding on cell surface. I.W. performed protein engineering and characterizing the binding kinetics. M.H. and J.L. performed three-color co-tracking. O.B. developed the orthogonal GFP binders and targets. U.R. contributed the HD10 anti-HaloTag nanobody. Y.M. produced the Wnt agonist under the supervision of K.C.G.. R.K. contributed to super-resolution imaging and image evaluations. C.Y. supervised the project. C.Y. and J.P. wrote the draft manuscript together with R.K. and contributions from all authors.

## Author ORCIDs

Changjiang You https://orcid.org/0000-0002-7839-6397 Jacob Piehler https://orcid.org/0000-0002-2143-2270 Rainer Kurre https://orcid.org/0000-0002-6872-6567

## Methods

### Materials

#### Production of dye-labeled nanobodies

Competent *E.coli* Rosetta (DE3) cells (Novagen) were transformed with the nanobody plasmids. Cells were grown at 37 °C in LB medium supplemented with 100 μg mL^−1^ ampicillin until an OD_600nm_ of 0.6–0.8 was reached. Protein expression was induced by adding 0.8 mM IPTG followed by overnight culturing at 18 °C. Cells were pelleted by centrifugation, resuspended in HEPES-buffered saline (HBS: 20 mM HEPES pH 7.5, 150 mM NaCl) and supplemented with DNAse, lysozyme and protease inhibitors. After lysis by ultracentrifugation at 55.000 × g, 25 min, 4 °C (Type 70 Ti, Beckman Coulter), the supernatant was purified by IMAC (5 mL HiTrap Chelating HP, GE Healthcare) using an FPLC system (ÄKTAprime, GE Healthcare). Proteins were eluted by a linear gradient against HBS buffer containing 500 mM imidazole. Collected nanobody fractions were then fractionated by size exclusion chromatography (SEC) using a Superdex 75 Increase 10/300 GL column (GE Healthcare) in HBS buffer.

Maleimide-cysteine reaction was used for site-specific fluorescence labeling of nanobodies. Maleimide-fluorophore conjugates used in the work were summarized in **Table S1**. Two-fold excess of maleimide-fluorophore conjugates diluted from 10 mM DMSO stocks was mixed with nanobodies for 30 min at room temperature. The reaction was stopped by addition of a 3-fold excess of free cysteine over the fluorophore for 15 min and further purified by SEC. Degree of labeling of all fluorophore-conjugated nanobodies was determined by UV/Vis spectroscopy using published (fluorescent dyes) or calculated (proteins) extinction coefficients and correction factors. Labeled nanobodies were flash-frozen in liquid nitrogen into aliquots and stored at -80 °C until use.

#### In *vitro* characterization of binding kinetics

The interaction kinetics of purified Nanobodies with different target was quantified by a home-built simultaneous real-time total internal reflection fluorescence spectroscopy and reflectance interference (TIRFS-RIF) detection in a flow-through system [64]. In brief, the setup employs white-light interference upon reflection at a 400 nm SiO_2_ layer on a glass transducer for label-free detection of protein binding. Simultaneous surface-sensitive fluorescence detection is enabled by laser excitation via total internal reflection using a glass prism. We used a 488 nm laser for excitation of mEGFP and fluorescence was filtered by bandpass filters between 495-605 nm for detecting by photomultipliers. Quantification of the binding kinetics of anti-AFLA nanobody mutant AnbAE is described below as example: for immobilization of His-tagged AnbAE (AnbAE-H6), a TIRFS-RIF transducer was coated with a dense PEG polymer brush that was functionalized with tris-(nitrilotriacetic acid) (tris-NTA) as described previously [65].

Protein interactions were probed under flow-through conditions using HBS. AnbAE-H6 was immobilized by His-tag on the Ni^2+^-loaded tris-NTA surfaces to yield low densities of ∼0.2-1 ng**·**mm^−2^. Tagless mEGFP-fused with ALFAtag in varying concentrations were injected for 60 s under a constant flow rate of 4.2 μL**·**s^−1^, followed with rinsing of HBS buffer with a flow rate of 10 μL**·**s^−1^. After each experiment, the surface was regenerated by washing with 500 mM imidazole in HBS. Kinetic and rate constants were extracted from fluorescence signal using the BIAevaluation 3.0 software (BIACORE) by applying a 1:1 Langmuir model.

#### Cell culture, transfection, live cell labeling and Wnt stimulation

The plasmids used for cell transfection are listed in **Table S2**. HeLa cells were cultivated at 37 °C under 5% CO_2_ in MEM with Earle’s salts supplemented with 10% FBS superior (Merck KGaA), 2 mM L-Alanyl-L-Glutamine (Biochrom), 1% non-essential amino acids (Merck KGaA) and 10 mM HEPES buffer (Carl Roth). Cells were transfected with single or multiple plasmids at 30-40% confluency by calcium phosphate precipitation overnight, followed by medium exchange and regeneration for 2-3 days. Before the day of microscopy imaging, cells were detached by room temperature treatment of Accutase (Innovative Cell Technologies) and seeded on microscopy cover slides coated with a 50/50 (w/w) mixture of poly-*L*-lysine-graft copolymers of polyethylene glycol (PLL-PEG) that were modified with an RGD-peptide (PLL-PEG-RGD) and a terminal methoxy group (PLL-PEG-OMe), respectively [66]. Meanwhile, 2 µM porcupine inhibitor IWP-2 (Sigma-Aldrich/Merck, I0536) was added to suppress secretion of endogenous Wnts. Labeling and microscopy imaging were performed in phenol red-free MEM medium supplemented with FBS. For nanobody labeling, 3 nM of dye-nanobody conjugate was added in the medium without further washing steps. Labeling of HaloTag and SNAP tag with dye-conjugates were carried out as described previously [46]. Briefly, 50 nM of dye-HaloTag Ligand (Promega, G8254 and G3221) or 20 nM of SNAP Surface 647 (New England Biolabs Inc., S9136S) were added to cell culture medium at 37 °C. After incubation for 20 min, cells were washed by warm PBS buffer for three times before refilling with phenol red-free MEM medium. For Wnt ligand stimulations, 50 nM recombinant carrier free human Wnt3a (R&D Systems Inc., 5036-WN-010/CF) and/or human Wnt5a (R&D Systems Inc., 645-WN-010/CF) were added to cell culture medium during the microscopy experiments. For stimulation by the Wnt surrogates [12, 13], 80 nM of the first-generation surrogate scFV-DKK1c (FGS) or 10 nM of the next generation surrogate (NGS) was added.

#### Microscopy sample preparation

Transfected cells were transferred to PLL-PEG-RGD-coated cover slides and kept in phenol-red free medium and incubated overnight (37°C, 5% CO_2_). In addition, cells transfected with Wnt receptors were treated with 2 µM of IWP-2 to avoid autonomous Wnt production. IWP-2 selectively blocks the activity of Porcupine, which is a multi-pass transmembrane *O*-acyltransferase in the ER responsible for Wnt palmitoylation. By this treatment, Wnt was retained in the ER and not secreted into the medium. The coverslip with cells was placed in an SDS/ethanol-cleaned home-built microscopy chamber for TIRF microscopy imaging.

#### Single-molecule TIRF microscopy

Single-molecule imaging was carried out by total internal reflection fluorescence microscopy (TIRFM) using an inverted microscope (IX83-P2ZF, Olympus) equipped with a motorized quad-line TIR illumination condenser (cellTIRF-4-Line, Olympus). The dyes Rho 11, Dy-647/SNAP Surface-649 and Dy-752 were excited using a 100× oil immersion objective (UPLAPO100XOHR, NA 1.5, Olympus) at 561 nm (2RU-VFL-P-500-560-B1R, MPB Communications), 642 nm (2RU-VFL-P-500-642-B1R, MPB Communications) and 730 nm (LuxX 730-50, max. 50 mW, Omicron), respectively. Laser lines were coupled into single-mode polarization-maintaining optical fibers which were connected to the 4-line TIR condenser. Laser power attenuation and blanking was achieved by a real-time controller (U-RTCE, Olympus). For the 561 and 642 nm laser, we used an acousto-optical tunable filter (TF525-250-6-3-GH18A, Gooch & Housego) linked to an eight-channel digital frequency synthesizer (MSD040-150-0.2ADM-A5H-8X1, Gooch & Housego). Fluorescence was filtered by a penta-band polychroic mirror (zt405/488/561/640/730rpc, Semrock) and excitation light was blocked by a penta-band bandpass emission filter (BrightLine HC 440/521/607/694/809, Semrock). Up to four channels could be simultaneously acquired by using the four quadrants of a single back-illuminated sCMOS camera (ORCA Fusion-BT, Hamamatsu) and a four-color image splitter (MultiSplit V2, CAIRN). The latter is equipped with three dichroic beamsplitters at 565 nm, 630 nm and 735 nm (T565LPXR, 630 DCXR and 735DCXR, Chroma) and four single-band bandpass emission filters (BrightLine HC 520/35, BrightLine HC 809/81, Semrock; ET 600/50, ET 685/50, Chroma). Time-lapse acquisitions at 30 frames per second were performed with a 2x2 binning and a final pixel size of 130 nm.

For dual channel imaging, simultaneous excitation at 560 nm and 642 nm lasers were employed in combination with dual-color image acquisition of the Rho11 (orange) and Dy647 (red) channel. For triple-color experiments, Rho11, Dy647 and Dy752 (dark-red) dyes were excited by the 561 nm, 642 nm and 730 nm laser lines simultaneously. For each channel, penetration depth of the evanescent field as well as laser excitation intensities (typically 50-500 W/cm^2^) were optimized to obtain comparable signal to background levels in each channel. Viable cells showing typical emitter densities of 0.1–0.8 copies/μm^2^ were imaged at 30 frames per second using CellSens 3.2 (Olympus) as acquisition software.

For long-term TALM imaging, 20,000 consecutive frames (11.1 min) were recorded. After a short pause (typically 1 min) for data storage, another time-lapse image stack of 20,000 frames was recorded. Five cycles of image acquisition were conducted to obtain continuous single molecule imaging for 1 hour. The focus was continuously stabilized during the experiment by a hardware autofocus-system (IX3-ZDC2, Olympus) using an internal laser diode at 830 nm. In all imaging experiments, an oxygen-scavenging system composed of glucose oxidase (4.5 U·mL^−1^), catalase (540 U·mL^−1^) and glucose (4.5 mg·mL^−1^) was added to increase photostability. Additionally, a photoprotectant redox system composed of ascorbic acid and methyl viologen (both 1 mM) was applied.

#### Single-molecule tracking

Single-, two- and three-color raw TIRF images were evaluated using an in-house developed Software for Localization-based Imaging in Matlab (SLIMfast) [42]. SLIMfast allows for single-molecule localization, rendering and tracking. Individual protein-protein interaction events were analyzed by single-molecule (co-)localization and (co-)tracking. Protein dynamics and confined diffusion behavior were determined by trajectory analysis based on step-length distributions and MSD analysis [43].

For channel registration, 200 nm TetraSpeck beads (Thermo Fisher Scientific) as multi-color fiducials visible in all fluorescence channels were used. While the TetraSpeck beads are not labeled with NIR dyes, the high brightness of the far-red channel can be used to obtain a reasonable crosstalk in the NIR-channel upon excitation at 642 nm. After bead localization in all spectral channels, we calculated projective transformation matrices to spatially align up to four channels with sub-pixel accuracy correcting for relative translation-, rotation- and scaling factors with respect to the defined reference channel.

Localization of individual fluorescence emitters against noise was done at a set error probability of 10^−5^ (less than 1 false positive detection per frame) with a point spread function (PSF) estimated from each respective channel using SLIMfast [67]. Single-molecule tracking was carried out using the algorithm u-track [68]. Upper boundaries for particle linking were established upon a prior robust evaluation of the frame-to-frame nearest-neighbor distribution. Gap closing with a maximum of 5 frames was allowed to account for missing localizations due to e.g. fluorescence blinking. Trajectories within an observation time (10 or 5 frames, see below) were used for further processing. Co-confinement analysis between spectral channels were carried out based on co-tracking. First, frame-by-frame co-localization within a radius of 150 nm led to co-localized species. Tracking of co-localized emitters by u-track with the same parameters as described above was performed. Molecules co-diffusing for ≥10 frames (≥ 350 ms) were identified as interaction events. Relative co-confinements were related to the least labeled receptor as it limits the absolute number of co-localization events:

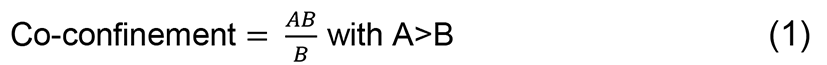

Here, A, B and AB are the total number of localizations observed for each individual receptor channel and the co-localized receptor subunits, respectively.

Diffusion constants were determined based on 150-frame TIRF image stacks acquired at video rate (5.2 s). For determining time-dependent change of diffusion constants (**Fig. 5F**, **Fig 6C**), the TIRF image stacks were chosen at each given time. Trajectories longer than 5 consecutive frames were used for analysis and step length (Δ𝑟) histograms based on the displacements of the 5 frames (Δ𝑡 = 175 ms) were computed. To estimate diffusion constants from the obtained histograms, we chose two-population model based on Bayesian information criterion with the diffusion constants 𝐷_1_ and 𝐷_2_ (slow and fast) and fitted the probability density function 𝑝(Δ𝑟, Δ𝑡) to the step length histograms:

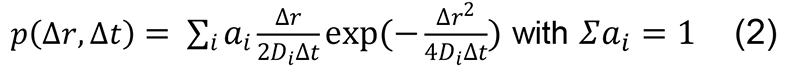

𝑎_𝑖_ is the proportions of the populations. The averaged diffusion constants were obtained by 𝐷 = ∑_𝑖_ 𝑎_𝑖_𝐷_𝑖_.

#### Degree of labeling (DOL)

The effective nanobody DOL on cell surface was determined by calculating the number of localized molecules in the nanobody channel [NB] and the SNAP channel [SNAP] acquired in the first 50 frames. In order to account for false-positive background localizations, untransfected HeLa cells were subjected to the exact same labeling procedure and image acquisition and used as the bg[NB] and bg[SNAP]. The calculation is based on the following equation:

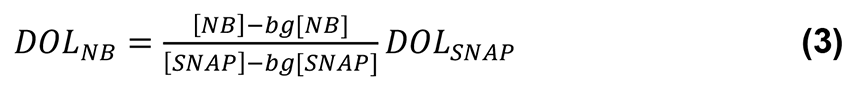

Here, 𝐷𝑂𝐿_𝑆𝑁𝐴𝑃_ is 43% according the calibration in HeLa cells [46].

#### Tracking and localization microscopy (TALM)

The principle of TALM is based on rendering single-molecule localizations maps with a defined observation window according to the diffusive behavior of the investigated protein species [43, 47]. Here, the localization maps were rendered with a final pixel size of 26 nm (one fifth of raw data pixel size) and a fixed global PSF radius of 26 nm (one pixel of localization map) for Gaussian blurring. As illustrated in **Fig. 3E-H**, time-lapse rbTALM images were rendered as TALM movies by moving a defined observation window with a certain time interval. Accumulated rbTALM images were reconstructed as time-lapse TALM but cumulating all localizations of the observation window until the given time. The obtained multi-color time-lapse TALM visualizes diffusion dynamics as well as local confinement. Co-localization TALM (coloc-TALM) images were reconstituted from the colocalized receptor pairs selected by frame-by-frame co-localization within a radius of 150 nm. The selection criterion obtained coloc-TALM images containing the same spatiotemporal information in each receptor channel (**Movie S7**). To illustrate different co-localized receptor pairs in one image, coloc-TALM images of the Lrp6, Fzd8, and Ror2 channel alone were shown to represent the co-localized Lrp6/Fzd8, Fzd8/Ror2 and Ror2/Lrp6, respectively.

Further quantitative analysis by pair correlation analysis of TALM images allows for quantification of spatiotemporal co-clustering and confinement zones [43, 47]. Based on previous simulations, unbiased pair correlation analysis requires each TALM image contains 100 localizations per µm^2^ [47]. Here, to explore the dynamics of nanoscale co-clustering, TALM movies were rendered with a moving observation window of 300-500 frames with a time interval of 280-350 ms (i.e. 1.4 – 2.9 min in total) based on the differences of localization density. In this way, the moving observation windows resulted in localization densities of > 100 localizations per µm^2^ for robust pair correlation analysis. For faithfully monitoring the time-course of Wnt signalosome formation, steps similar or equal to the timespan of observation window were applied. Particularly, stepwise of 1.75 min for visualizing the formation of canonical Wnt signalosome (**Fig. 4**), 1.4 min for the common Wnt signalosome (**Fig. 5**), 2.1 min for the noncanonical Wnt signalosome (**Fig. 6**) was used and specified in each figure.

For pair correlation analysis of TALM images, the time interval of more than 8 frames (280 ms) was chosen to reliably investigate co-clustering and confinement zone formation with strong reduction of the diffusion dependent false-positive confinements. If the time interval is too short, those false-positives could result in false localization clusters. Previous simulations had shown the minimum interval for unbiased pair correlation analysis of a freely diffusing receptor was 256 ms for *D* = 0.05 µm^2^/s and 160 ms for *D =* 0.1 µm^2^/s [47]. We therefore chose time intervals 280 ms as the average diffusion constants for Ror2, Fzd8 and Lrp6 in the resting state of three-color rbTALM experiments were determined in the range of 0.18 to 0.05 µm^2^/s.

#### Time-lapse pair correlation analysis of TALM images

Pair correlation analysis of TALM images was carried out as previously described [47]. First, a conventional image correlation spectroscopy (ICS) analysis was applied to TALM images yielding the pair correlation function *g(r)* in polar coordinates which is fitted by a two-component model [50]:

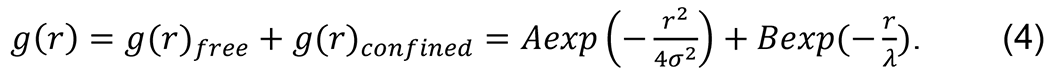

Here, *A* is the amplitude of free diffusion, σ is the standard deviation (i.e., the average single-molecule localization precision, σ = 26 nm in this work), *B* is the amplitude of confined diffusion, and λ is the average radius of the randomly shaped confinement zones. The confinement probability (𝑐𝑝) was defined to describe the chance of the diffusive species being confined in the analyzed region. For co-confined diffusion of receptor pairs, 𝑐𝑐𝑝 was used to denote the co-confinement probability. Both are calculated as:

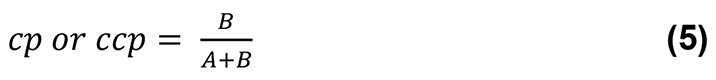

By implementing pair correlation analysis for consecutive images within TALM stacks, the obtained 𝑐𝑝 and 𝑐𝑐𝑝 values of each TALM image were plotted against time of the TALM images. Mono-exponential fitting was applied to the time-dependent 𝑐𝑝 and 𝑐𝑐𝑝 to obtain the characteristic receptor assembly time (*t_a_*) as the 1/e value.

Since co-clustering of the Wnt co-receptors was most pronounced at cell periphery (**Fig. S8A, E**), we selected region-of-interests (ROIs) at the cell rim for analysis. Dimensions of the ROIs were 5x5 µm^2^ which were the typical sizes used in previous pair correlation analyses [47, 50]. To further enhance the robustness of quantification, three representative ROIs were analyzed and the results were reported as the mean value ± standard deviation.

## Statistics

Statistics of histograms or box charts in **Fig. 1J-M**, **Fig. 2E, F**, **Fig. 5D** were calculated as two sample t-test with sample volumes reported in each figure. Histograms in **Fig. 1E, F, H, I**; **Fig. S1E, F**; **Fig. S2C-F**; **Fig. S3D, H, L**; **Fig. S5L**; **Fig. S6C-F**; **Fig. S9D-F**; **Fig. S11E, G** were fitted using Matlab or Origin by Gaussian mixture models and 1 or 2 populations as indicated in each figure. Standard deviations are based on Gaussian fits with 95 % confidence interval. Exponential fittings in **Fig. 4J, M**, **Fig. 5 J, L, K**, **Fig. 6D**, **Fig. S4E, G** with the mean ± standard deviations (σ) and replicates were indicated in each figure.

## Data availability

The datasets generated during and/or analyzed during the current study as well as microscope configuration files are available from the corresponding authors on reasonable request.

## Software and resources

The software SLIMfast for image analysis were previously programmed in-house [42]. SLIMfast requires Matlab version 2013b (Mathworks). The pair correlation analysis tool is another in-house developed Matlab script [47] which can be run in Matlab version after 2013b. Plasmids and software used in this study are summarized in supplementary Table S2, 3 which are available via the lead contact upon request. Further information and requests for resources and reagents should be directed to and will be fulfilled by the Lead Contact: Changjiang You.

## SI Figures

**Fig. S1.**
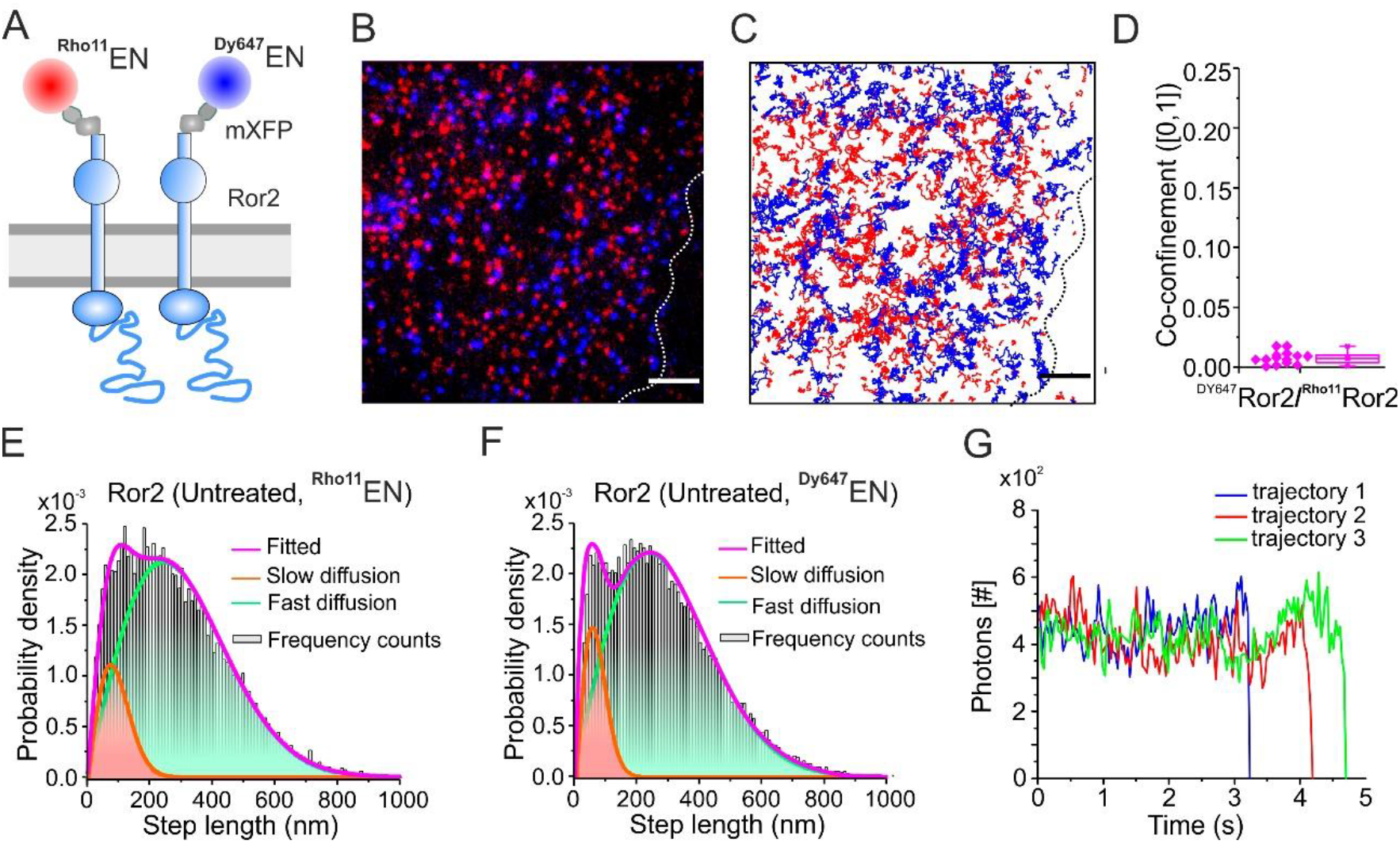
Two-color single-molecule tracking assays of Ror2 pre-homodimerization. (A) Cartoon of labeling mXFPe-Ror2 by anti-GFP enhancer nanobodies (EN) conjugated with Dy647 and Rho11. (B) Representative two-color raw TIRF microscopy image for cells at the resting state. (C) Single-molecule trajectories of ^Dy647^EN (blue) and ^Rho11^EN (red)-labeled mXFPe-Ror2 in a representative cell. Scale bars (B, C): 5 µm. (D) Co-confinement of two-color labeled Ror2. (E, F) Step length histogram analysis for the diffusion of ^Rho11^EN-labeled Ror2 (E) and ^Dy647^EN-labeled Ror2 (F), respectively. Number of steps in histogram (E) 29448 and (F) 18136. Nonlinear least square fit with two fractions (*i.e.* as the fast and slow diffusion) was applied to the step length histogram (gray bars). Fitted probability (magenta curve) consist of the fast (green) and the slow (orange) portions. (G) Photons vs time of the tracked individual ^Dy647^EN-labeled Ror2 show single photobleaching step.

**Fig. S2.**
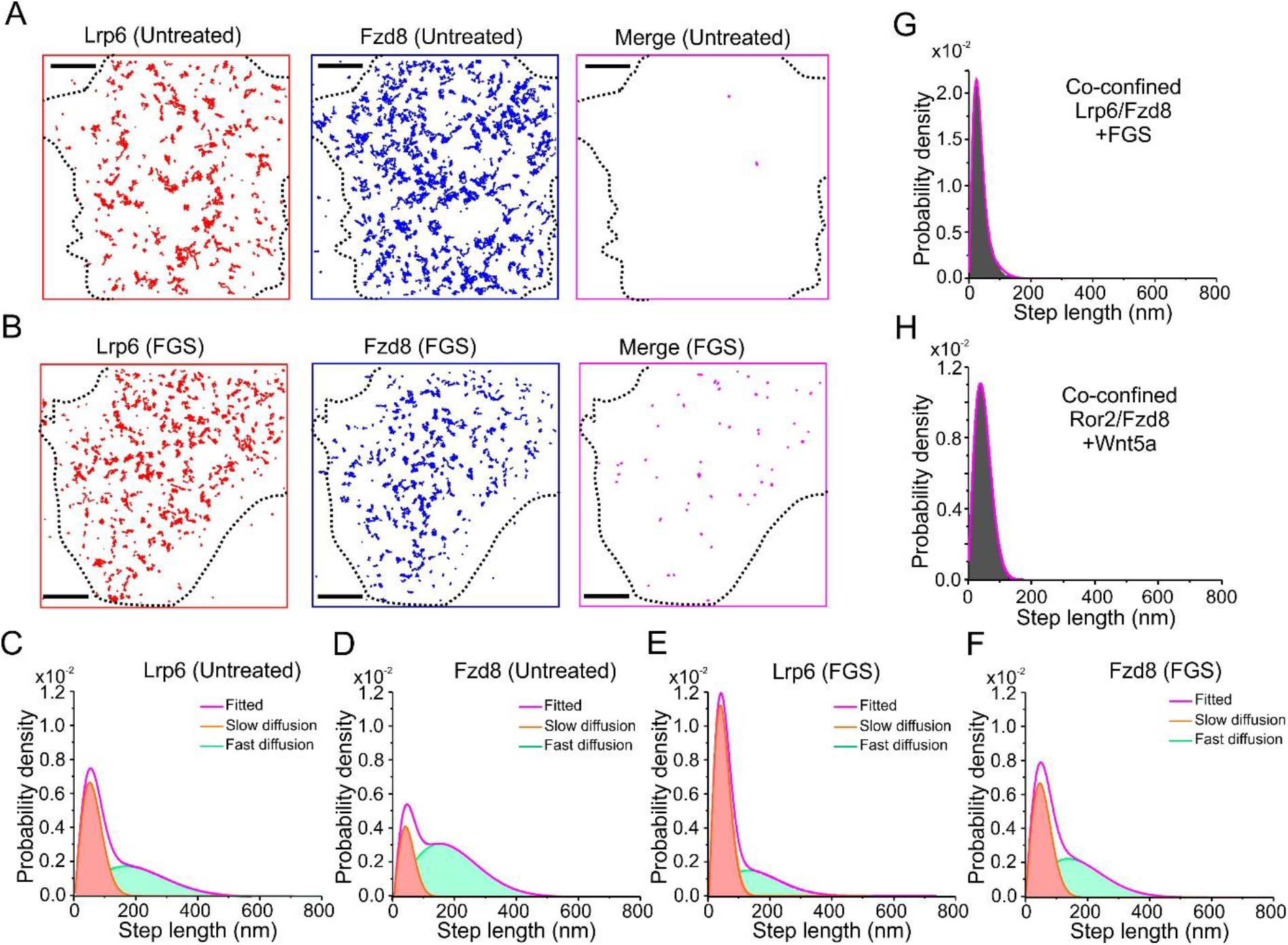
Two-color single-molecule tracking of Lrp6/Fzd8 stably labeled via EN and MI nanobodies. (A) Single-molecule trajectories of Lrp6 (red) and Fzd8 (blue) in the untreated state show minimum background co-confinement (magenta). Scale bars: 5 µm. (B) Trajectories of nanobody-labeled Lrp6 (red) and Fzd8 (blue) after stimulation by Wnt3a surrogate show significant slowing down and receptor co-confinement (magenta). Scale bars: 5 µm. (C, D) Step length histogram of Lrp6 (C) and Fzd8 (D) in the untreated state. Number of steps in histogram (C): 27686 and (D): 17423. (E, F) Step length histogram of Lrp6 (E) and Fzd8 (F) after canonical Wnt surrogate FGS stimulation, showing the increase of slow diffusion and decrease of fast diffusion. Number of steps in histogram (E): 46079 and (F): 25473. (G) Step length histogram of co-confined Lrp6/Fzd8 after stimulation by FGS. Number of steps: 2249. (H) Step length histogram of co-confined Ror2/Fzd8 after stimulation by Wnt5a (cf. Fig 1G). Number of steps: 949. The majority of co-confined receptors in panel (G) and (H) is slow diffusion.

**Fig. S3.**
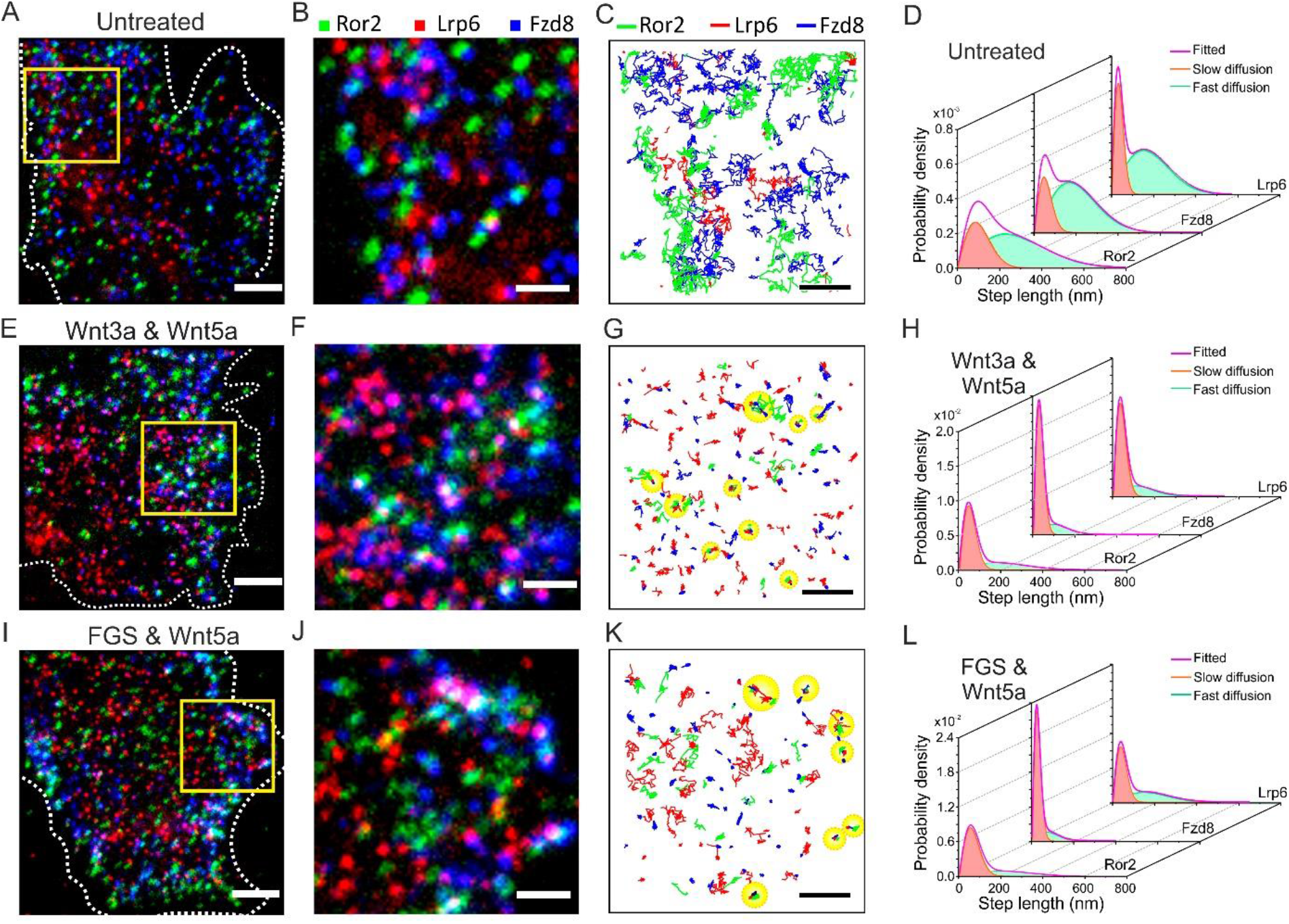
Three-color single-molecule tracking of Lrp6/Fzd8/Ror2. (A) Representative TIRF microscopy image of Ror2 (green), Lrp6 (red) and Fzd8 (blue) before stimulation, i.e. at untreated state. (B) Zoom-up of the highlighted region of (A). (C) Single-molecule trajectories of Ror2, Lrp6 and Fzd8 of the zoom-up region. (D) Step length histogram and fitting for the diffusion of each receptor at the untreated state. (E) Raw TIRF microscopy image of Ror2 (green), Lrp6 (red) and Fzd8 (blue) after adding the mixture of Wnt3a and Wnt5a. (F) Zoom-up of the highlighted region in panel (E). (G) Single-molecule trajectories of Ror2, Lrp6 and Fzd8 of the zoom-up region after stimulation by Wnt3a and Wnt5a. Co-confinement is marked in yellow. (H) Step length histogram and fitting for the diffusion of receptors by simultaneous Wnt3a and Wnt5a stimulation. (I) Raw TIRF microscopy image of Ror2 (green), Lrp6 (red) and Fzd8 (blue) after adding the mixture of Wnt3a surrogate FGS and Wnt5a. (J) Zoom-up of the highlighted region in panle (I). (K) Single-molecule trajectories of Ror2, Lrp6 and Fzd8 of the zoom-up region. Co-confinement is marked in yellow. (L) Step length histogram and fitting for the diffusion of receptors by simultaneous stimulation with FGS and Wnt5a. Scale bars (A, E, G): 5 µm. Scale bars (B, C, F, G, J, K): 2 µm.

**Fig. S4.**
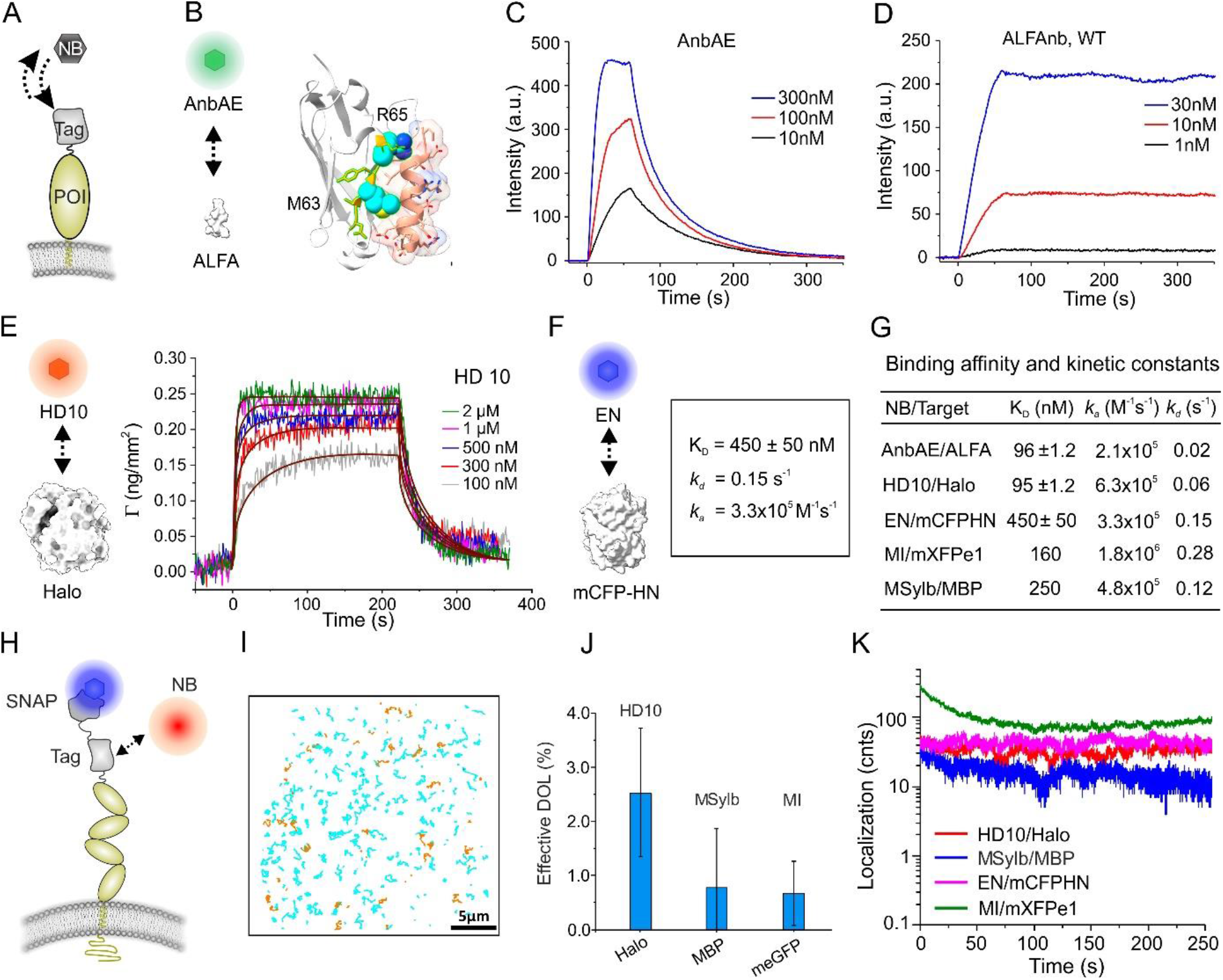
Engineered nanobodies for long-term single molecule localization microscopy. (A) Scheme illustrating reversible binding of nanobody to transmembrane receptor. (B) Designed point mutations of ALFA nanobody (AnbAE) for reverse binding with ALFAtag. (C) Quantification of the binding kinetics of ALFA-tag GFP to immobilized AnbAE by surface-sensitive total internal reflection fluorescence spectroscopy (TIRFS). Curves show ALFA-tag GFP added at different concentrations. (D) Stable binding of ALFA-tag GFP to ALFA nanobody wild type as comparison. (E) Reflectance interference spectroscopy (RIFS) of anti-Halo tag nanobody HD10 binding to immobilized HaloTag. Global fitting of binding and dissociation rate constants were extracted by applying a 1:1 Langmuir model to the curves of different HD concentrations (brown lines). (F) Reverse binding of antiGFP nanobody enhancer to mCFP*HN*. Binding constants were obtained by TIRFs-RIFs measurement. (G) Summary of nanobody binding affinity and kinetic constants determined in *vitro* by TIRFs-RIFs. (H) Scheme for quantification of the effective degree-of-labeling (*DOL*) at cell surface. (I) Two-color single-molecule tracking trajectories of chimeric SNAP-Halo transmembrane receptor model in living cell. Cyan: ^Dy647^SNAP. Orange: ^Rho11^HD10. Brown: Colocomotions. Scale bar: 5µm. (J) Effective *DOL* of different nanobodies determined at cell surface. (K) Plots of localization counts versus time for different combinations of nanobody/target. Localizations were obtained from TIRF microscopy acquired at video rate. PDB entries for structures in panel (B, E, F): 6I2G (ALFA tag and nanobody); 5Y2Y (HaloTag); 3K1K (green fluorescent protein bound to enhancer nanobody).

**Fig. S5.**
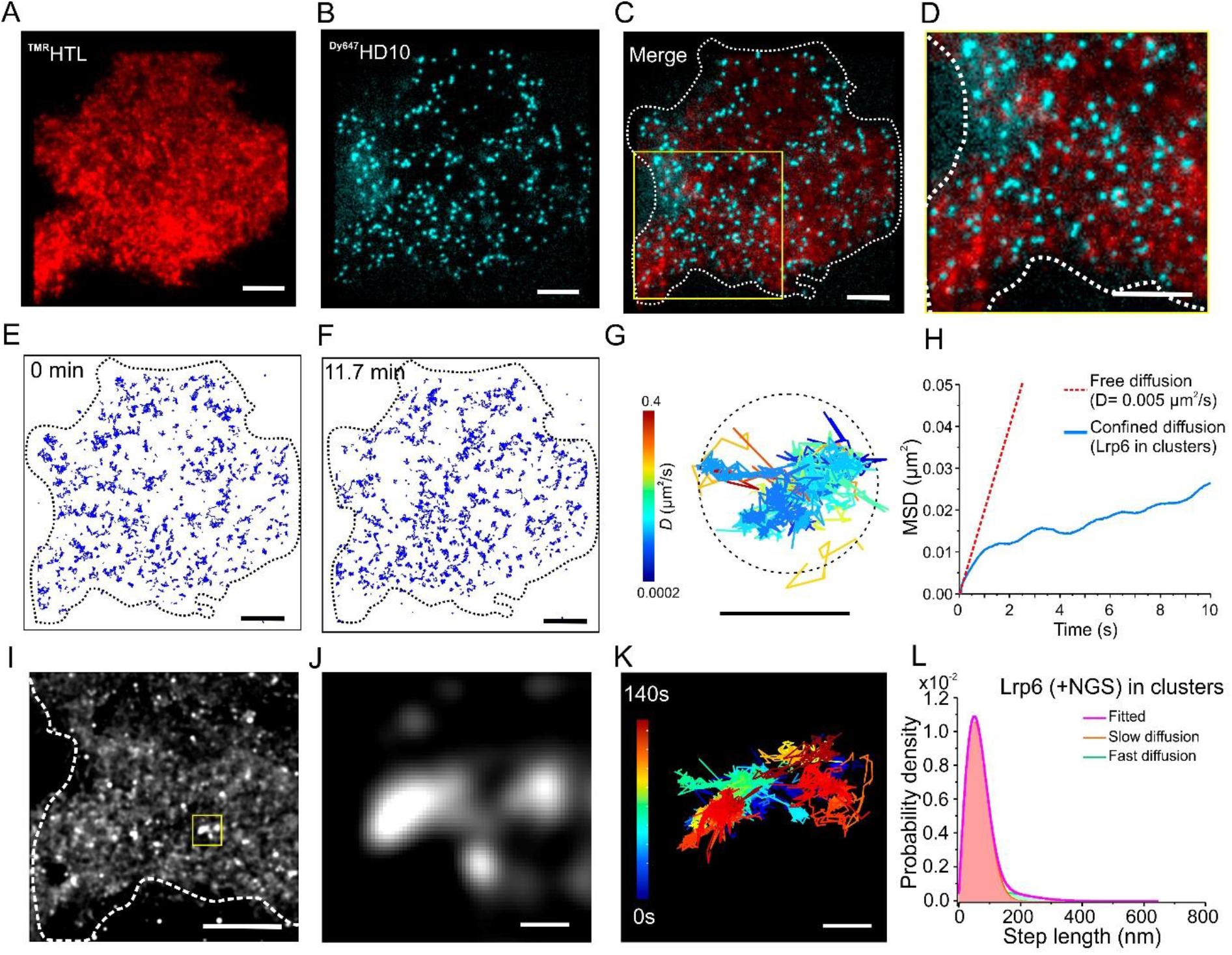
Comparison of covalent labeling and reversible nanobody labeling of HaloTag-Lrp6 in live cell. (A-C) Raw two-color TIRF microscopy image of HaloTag-Lrp6 labeled by TMR-HTL (A, red) and ^Dy647^HD10 nanobody (B, Turquoise) and the merged image (C) at time zero. Dashed line indicates cell boundary. (D) Zoom-up region of the square marked in panel (C). Scale bars (A-D): 5 µm. (E-F) Single molecule trajectories of ^Dy647^HD10-labeled Lrp6 recorded at the start (E) and the end of experiment (F). Scale bars: 5 µm. (G) Trajectories of ^Dy647^HD10-labeled Lrp6 in the confined diffusion region (dash circle) recorded for 84 s. Color-coded by diffusion constant (logarithmic scale). Scale bar: 500 nm. (H) Mean square displacement (MSD) analysis of a trajectory in the confined diffusion region (blue curve). Red dash line is the MSD plot of free diffusion for comparison. (I) Overview TALM image of the marked square in panel (C) obtained by accumulating the localizations of ^Dy647^HD10-labeled Lrp6 through the experiment. Scale bar: 500 nm. (J) Lrp6 clustering in the ROI highlighted in (I). (K) Trajectories of ^Dy647^HD10-labeled Lrp6 in the clustering region recorded for 140 s. Scale bars (J, K): 200 nm. (L) Step-length histogram of the trajectories in panel (K), showing confined diffusion of Lrp6 in the clusters.

**Fig. S6.**
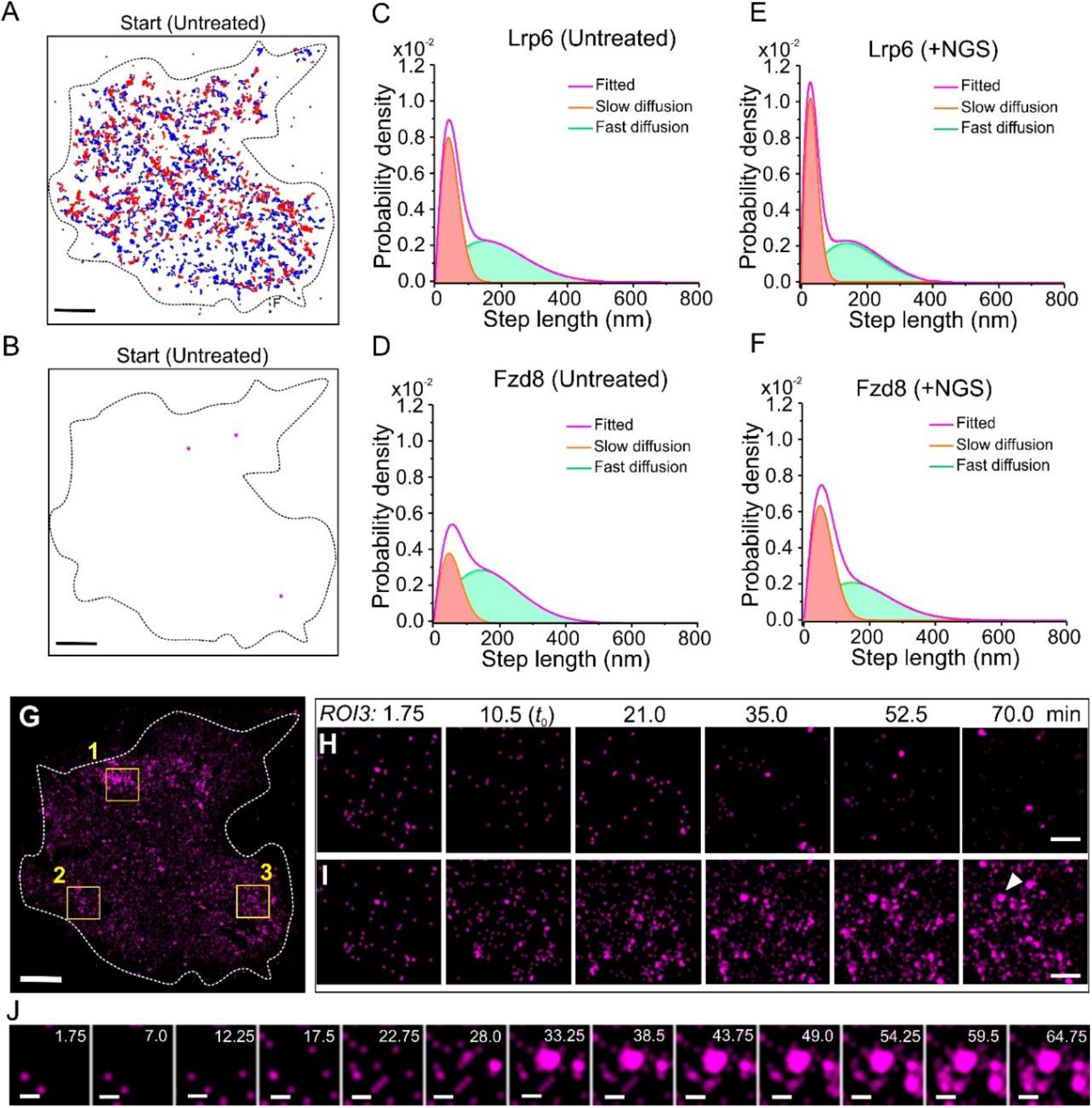
Two-*color* single-molecule tracking and rbTALM of Lrp6/Fzd8. (A) Single molecule trajectories of the reversibly ^Rho11^HD10-labeled HaloTag-Lrp6 and ^Dy647^EN-labeled CFP*HN*-Fzd8 at the untreated state. (B) Co-diffusion trajectories of Lrp6/Fzd8 for 10 steps at the untreated state. Scale bars: 5 µm. (C, D) Step length histogram analyses of Lrp6 (C) and Fzd8 (D) diffusion in the untreated state. (E, F) Step length histogram analyses of Lrp6 (E) and Fzd8 (F) in the same cell after stimulation with NGS for 60 min. (G) Overview rbTALM image of colocalized Lrp6 with Fzd8 (Lrp6/Fzd8, magenta) for a 70 min experiment. Altogether 12,000 frames with an interval of 350 ms were used which were retrieved from 120,000 frames raw data (35 ms per frame). Scale bar: 5 µm. (H, I) Time-lapse TALM images (H) and accumulated TALM images (I) of ROI3. Scale bars: 1 µm. Montages show the colocalized Lrp6/Fzd8 (magenta) in a 500-frames observation window (2.92 min) sliding at a stepwise of 300 frames (1.75 min). *t* is the step time. (J) Zoomed TALM image of Lrp6/Fzd8 co-clusters marked by white triangle in panel (I). Scale bars: 200 nm.

**Fig. S7.**
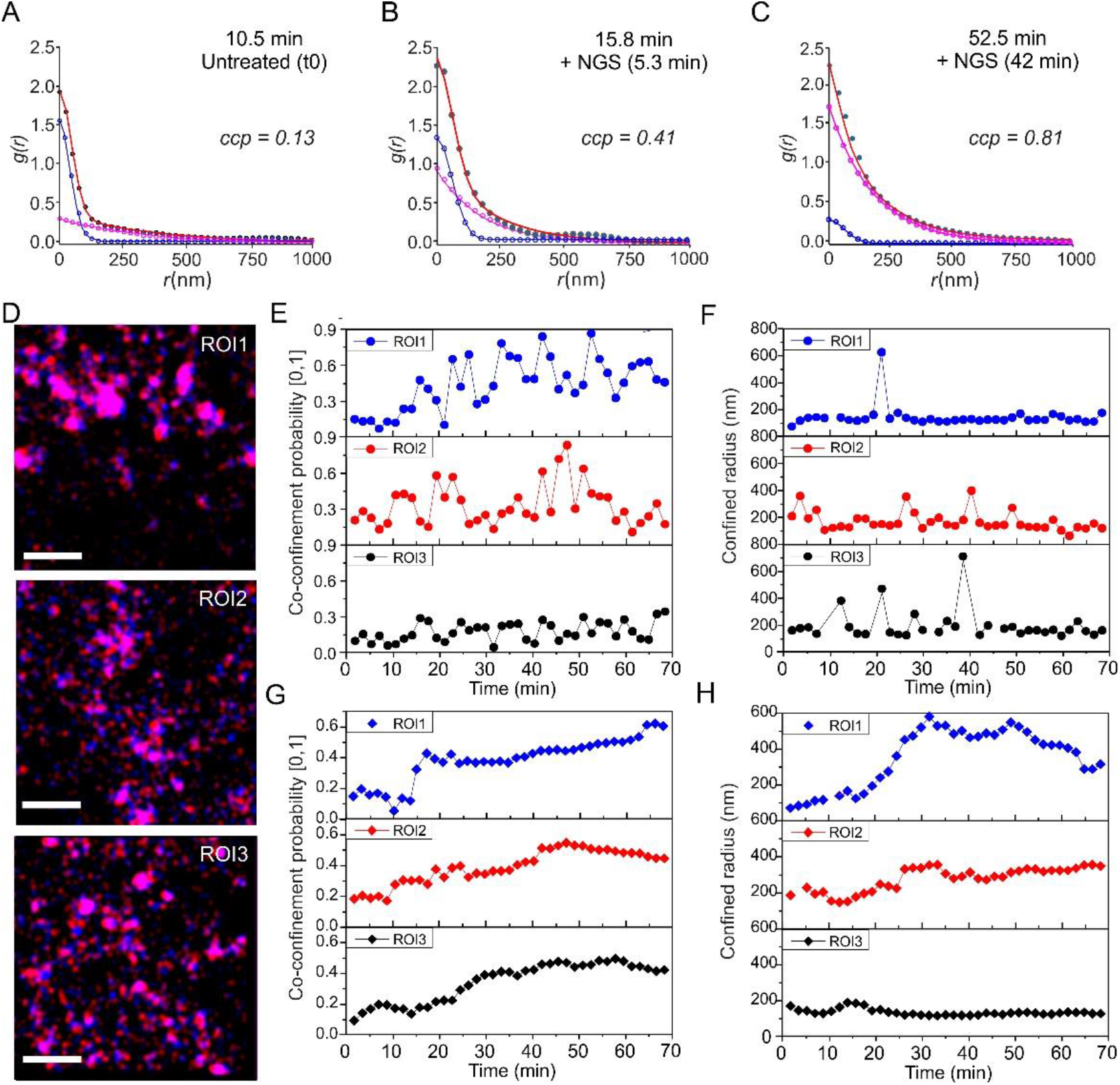
Pair correlation analyses two-color colocalization TALM (coloc-TALM) of canonical Wnt signalosome. (A) Pair correlation analysis curves of time-lapse TALM images in ROI1 at 8.75 min, i.e., 1.75 min before t_0_ of adding NGS. Co-confinement probability (*ccp*) is 0.13. Confinement radius (*λ*) is 123 nm. (B) Pair correlation analysis curves of ROI1 at 15.75 min (5.25 min after adding NGS), *ccp* = 0.41. *λ* = 114 nm. (C) Pair correlation analysis curves of ROI1 at 52.5 min (42 min after adding NGS). *ccp* = 0.81. *λ* = 116 nm. (D) Zoomed coloc-TALM images of the ROI1, 2 and 3. Colocalized Lrp6 (red) and Fzd8 (blue) are shown separately. Their overlapping is shown as magenta Scale bars: 1 µm. (E, F) Pair correlation analysis of time-lapse TALM for quantifying the kinetics of Lrp6/Fzd8 co-clustering upon NGS stimulation. Co-confinement probabilities (E) and confined radii (F) of each ROIs are plotted in column. (G, H) Pair correlation analysis of accumulated TALM images in three ROIs to obtain the average co-confinement probabilities (G) and confined radii (H).

**Fig. S8.**
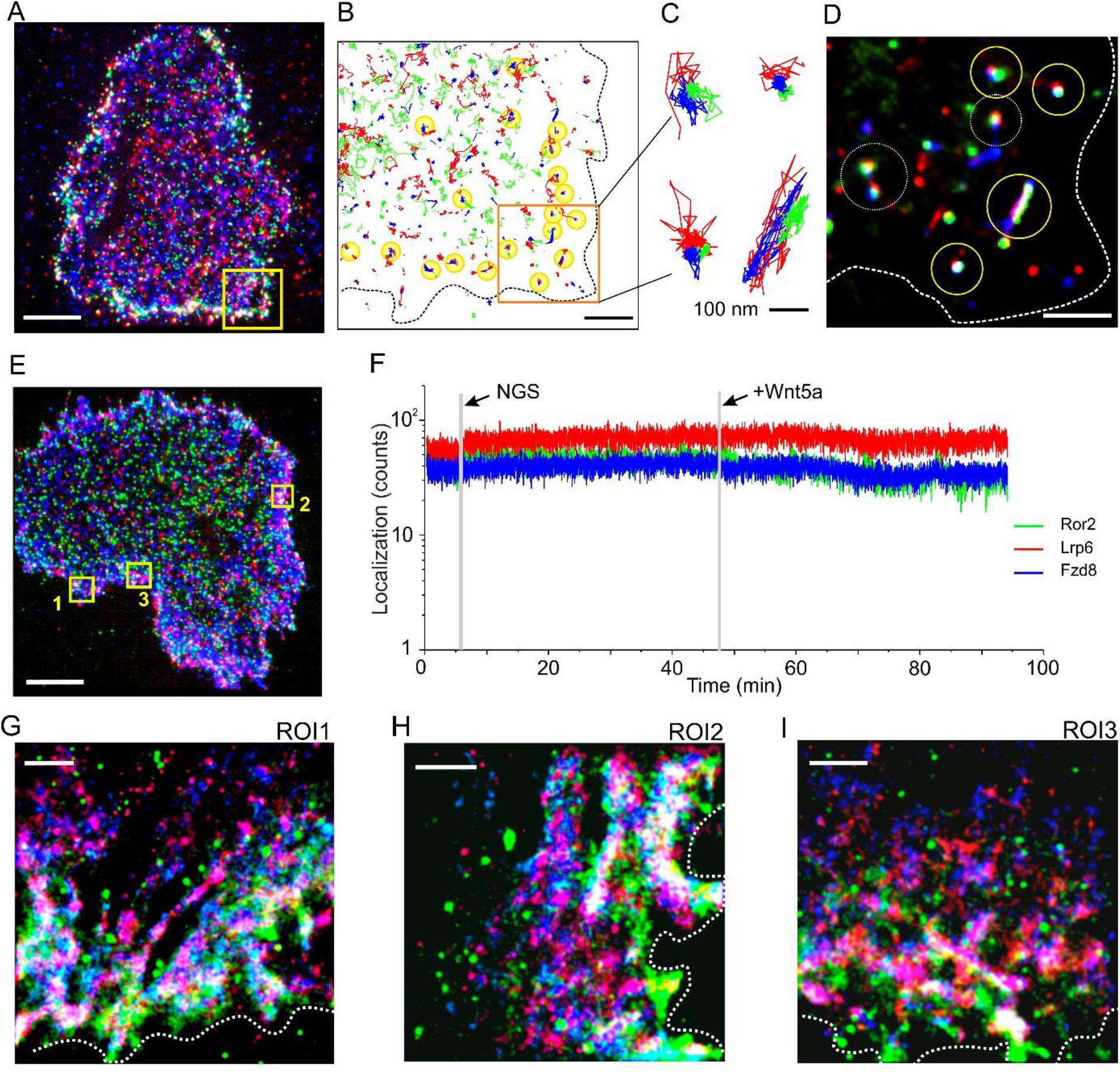
Three-color rbTALM imaging of the common Wnt signalosome. (A) Raw TIRF microscopy image of ^AnbAE^Ror2 (green), ^HD10^Lrp6 (red) and ^EN^Fzd8 (blue) in a live cell. Scale bar: 10 µm. (B) Trajectories in the marked yellow square, see also Fig. 5B. Scale bar: 2 µm. (C) Highlighted trajectory overlapping in the orange square based on 300-frame tracking. (D) rbTALM image of the orange square based on the 300-frame tracking. Scale bar: 1 µm. (E) Raw TIRF microscopy image of another live cell. Scale bar: 10 µm. ROIs are marked by numbered yellow squares and shown in Fig. 5 and Fig. S9, 10. (F) Plot of receptor localizations *vs* time as sum of the three ROIs. (G-H) Zoomed-up three-color rbTALM images of the ROI1 (G), ROI2 (H) and ROI3 (I) shown in panel (E). The accumulated three-color rbTALM images are based on accumulating 20105 frames with an interval of 280 ms which were retrieved from 160,840 frames acquired in 93.8 min. Ror2 (green), Lrp6 (red) and Fzd8 (blue). Scale bars: 1 µm.

**Fig. S9.**
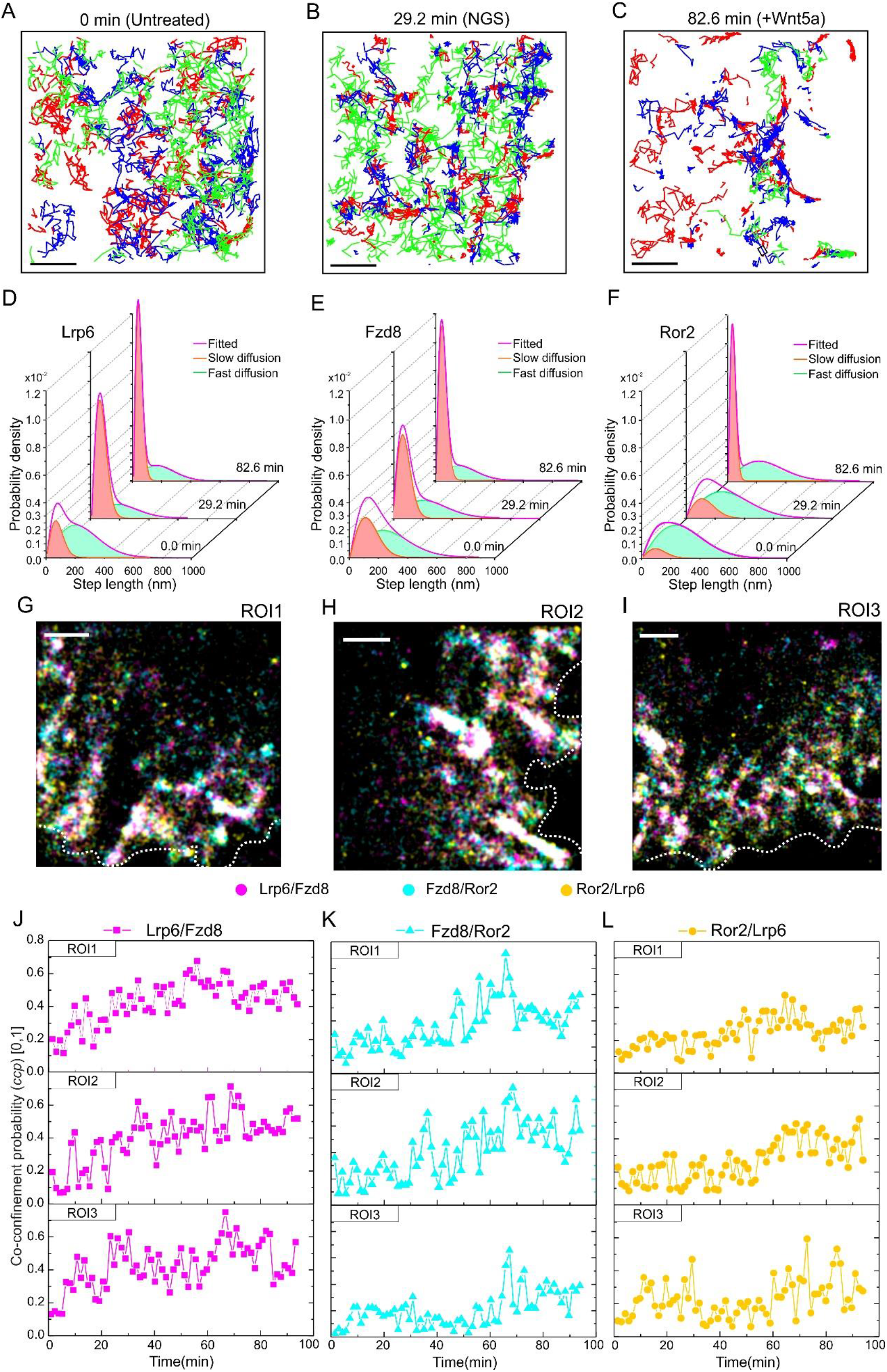
Time-dependent diffusion constant and time-lapse pair correlation analyses of the common Wnt signalosome. (A-C) Trajectories of Ror2 (green), Fzd8 (blue) and Lrp6 (red) in ROI2 recorded at different time and stimulation states. (D-F) Step length histogram analysis for determining the diffusion constants of Ror2 (D), Fzd8 (E) and Lrp6 (F) at 0 min of the untreated state, at 29.2 min by NGS stimulation and at 82.6 min with simultaneous stimulation of NGS and Wnt5a. (G-I) Accumulated three-color colocalization TALM images for ROI1 (G), ROI2 (H) and ROI3 (I) of the entire experimental time (93.8 min). Scale bars: 1 µm. Co-localized Lrp6/FZd8 (magenta), Fzd8/Ror2 (cyan), and Ror2/Lrp6 (orange) are shown with distinct colors. (J-L) Plots of co-confinement probability (*ccp*) *vs* time for the three ROIs. (J) Lrp6/Fzd8 (magenta square). (K) Fzd8/Ror2 (cyan triangle). (L) Ror2/Lrp6 (yellow dot). *ccp* values were quantified by pair correlation analysis of the time-lapse coloc-TALM images, which were obtained by accumulating colocalized receptors in a 300-frame observation window (1.4 min) and sliding at a stepwise of 300-frame (1.4 min). Frame interval is 280 ms.

**Fig. S10.**
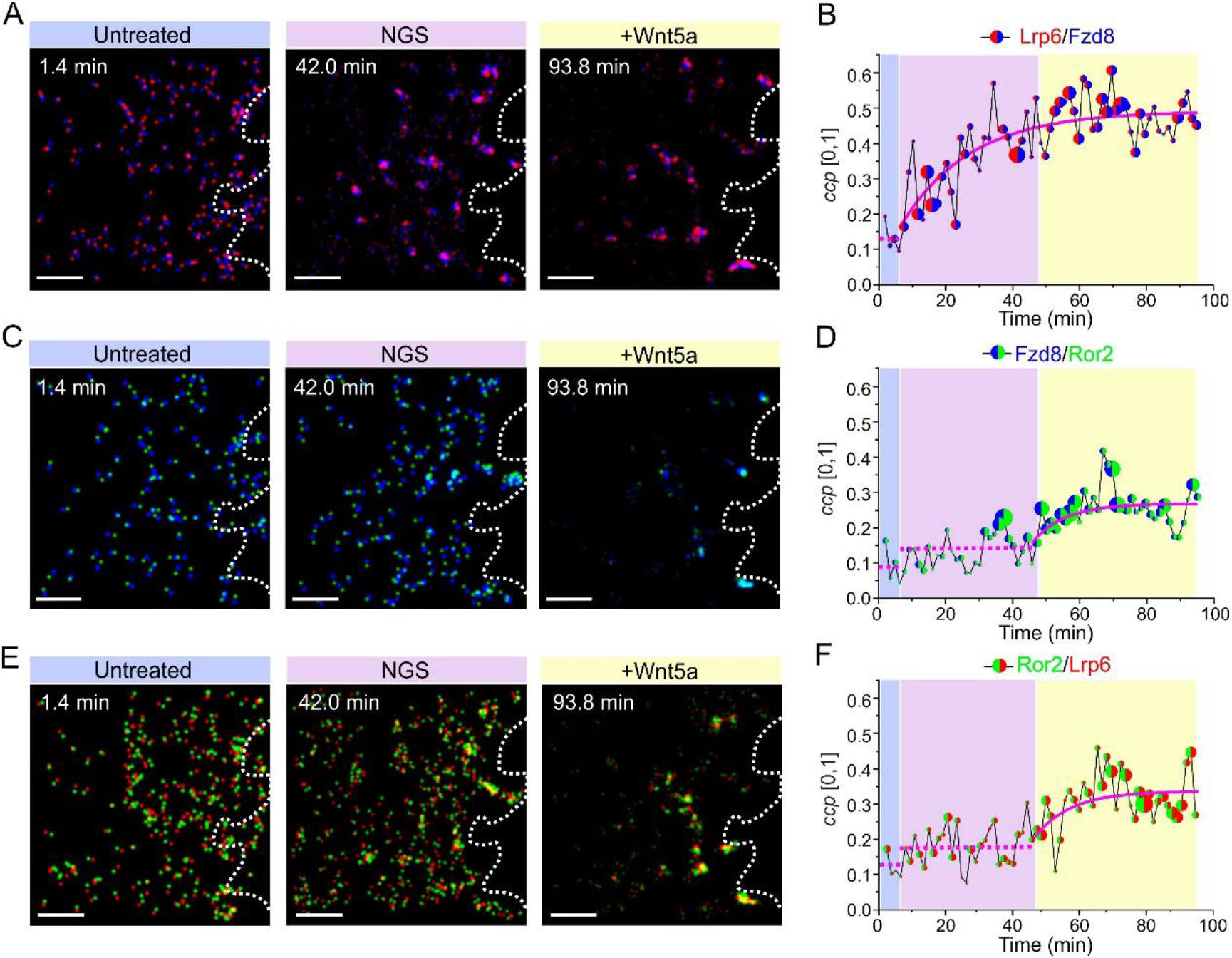
Kinetics of receptor co-clustering quantified by time-lapse two-color coloc-TALM and pair correlation analyses. (A) Two-color time-lapse coloc-TALM images of co-localized Lrp6 (red)/Fzd8 (blue) in ROI2 at given time by accumulating localizations in a 1.4 min observation window and sliding with a stepwise of 1.4 min. Receptor overlapping is shown in magenta. Untreated state (aqua blue), NGS stimulation (pink) and NGS+Wnt5a stimulation (yellow) are marked with color bars. (B) Co-confinement probabilities (*ccp*) for Lrp6/Fzd8 plotted as the mean values of the three ROIs marked in FigS9. Bubble size in Pair correlation analysis quantification curve represents the standard deviation. (C) Two-color time-lapse coloc-TALM images of co-localized Fzd8 (blue)/Ror2 (green) in ROI2. Receptor overlapping is shown in cyan. (D) Pair correlation analysis quantification of co-confinement probabilities of Fzd8/Ror2 in three ROIs. (E) Two-color time-lapse coloc-TALM images of co-localized Ror2 (green)/Lrp6 (red) in ROI2. Receptor overlapping is shown in yellow. (F) Pair correlation analysis quantification of co-confinement probabilities of Ror2/Lrp6 from three ROIs. Scale bars: 1 µm. Mono-exponential fittings are shown as magenta lines. Dash magenta lines are the mean values within the timespan used for eye-guidance as the fitting is not applicable.

**Fig. S11.**
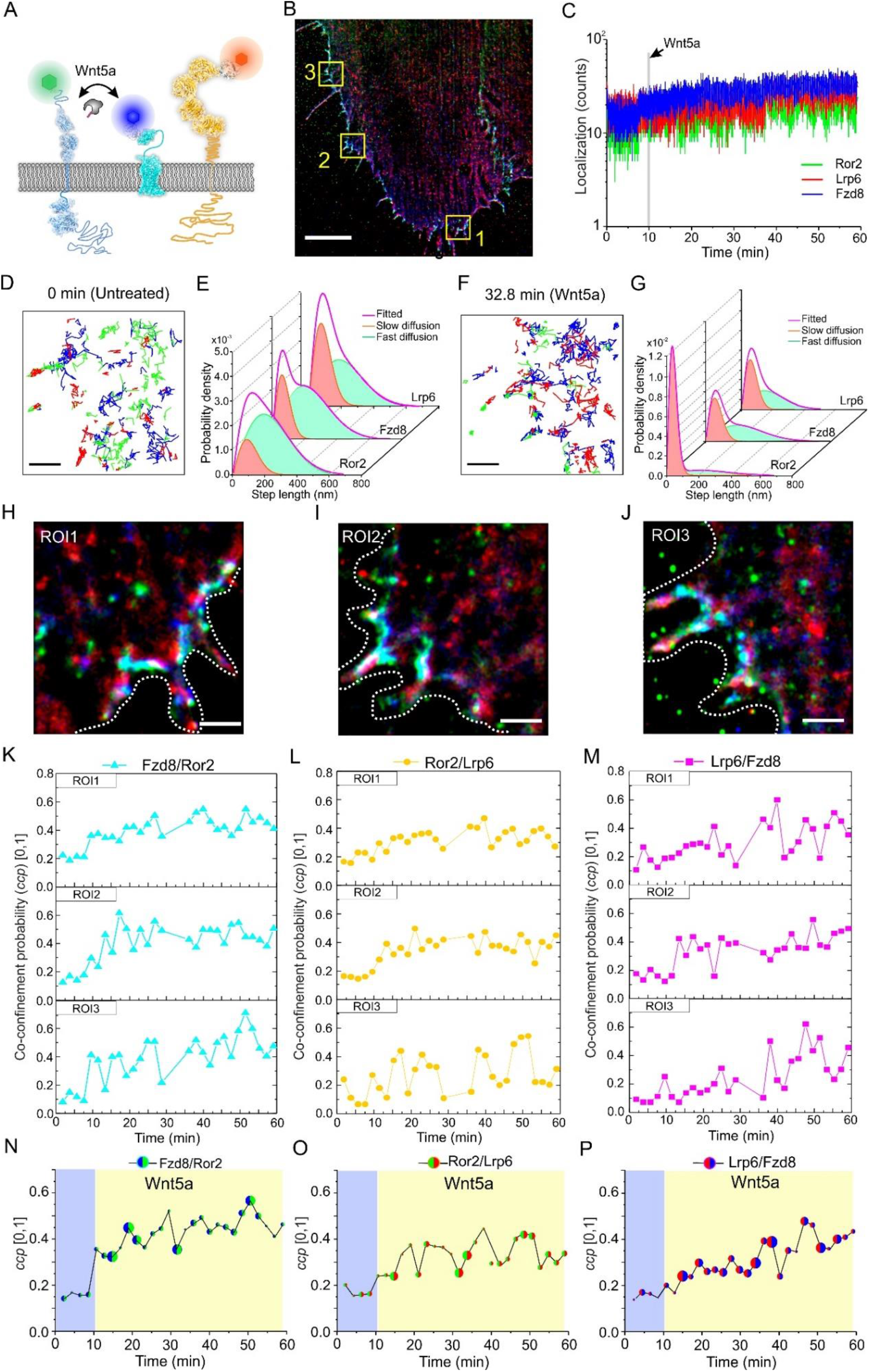
Wnt5a-induced Ror2/Fzd8 assembly in absence of canonical Wnt ligand stimulation. (A) Scheme of activating noncanonical Wnt signaling by Wnt5a in cells transiently expressing ALFA-Ror2, CFP*HN*-Fzd8, HaloTag-Lrp6 and orthogonally labeled with nanobodies. Wnt5a which dimerizes Ror2 and Fzd8 was added to induce noncanonical Wnt signaling. (B) Overview rbTALM obtained by accumulating 12,500 frames with an interval of 280 ms after adding Wnt5a, which were retrieved from 100,000 frames acquired in 58.3 min. Ror2 (green), Lrp6 (red) and Fzd8 (blue). Scale bar: 10 µm. (C) Plots of single molecule localization of receptors *vs* time. Localizations are sum of three ROIs marked in panel (B). (D) Trajectories of Ror2 (green), Fzd8 (blue) and Lrp6 (red) in ROI3 recorded at 0 min of the experiment, i.e., the untreated state. (E) Step length histogram analysis for determining diffusion constants of Ror2, Fzd8 and Lrp6 in the untreated state. (F) Trajectories of Ror2 (green), Fzd8 (blue) and Lrp6 (red) in ROI3 recorded at 32.8 min of experiment, i.e., by Wnt5a stimulation. (G) Step length histogram analysis for determining diffusion constants of Ror2, Fzd8 and Lrp6 by Wnt5a stimulation. (H-J) Accumulated three-color rbTALM of noncanonical Wnt signalosome including Ror2 (green), Lrp6 (red) and Fzd8 (blue). Zoom-ups of ROI1 (H), ROI2 (I) and ROI3 (J) marked in panel (B). Scale bars: 1 µm. (K, L, M) Plots of co-confinement probability (*ccp*) *vs* time for receptor co-clustering in the ROIs. (K) Fzd8/Ror2 (cyan triangle). (L) Ror2/Lrp6 (yellow dot). (M) Lrp6/Fzd8 (magenta square). Time-lapse coloc-TALM images were obtained by accumulating co-localized receptors in a 2.1 min observation window (450 frames, 280 ms interval) sliding with a stepwise of 2.1 min. (N, O, P) Time-dependent *ccp* of Fzd8/Ror2 (N), Ror2/Lrp6 (O) and Lrp6/Fzd8 (P) obtained as mean of three ROIs. Bubble size represents the standard deviation.

**Fig. S12.**
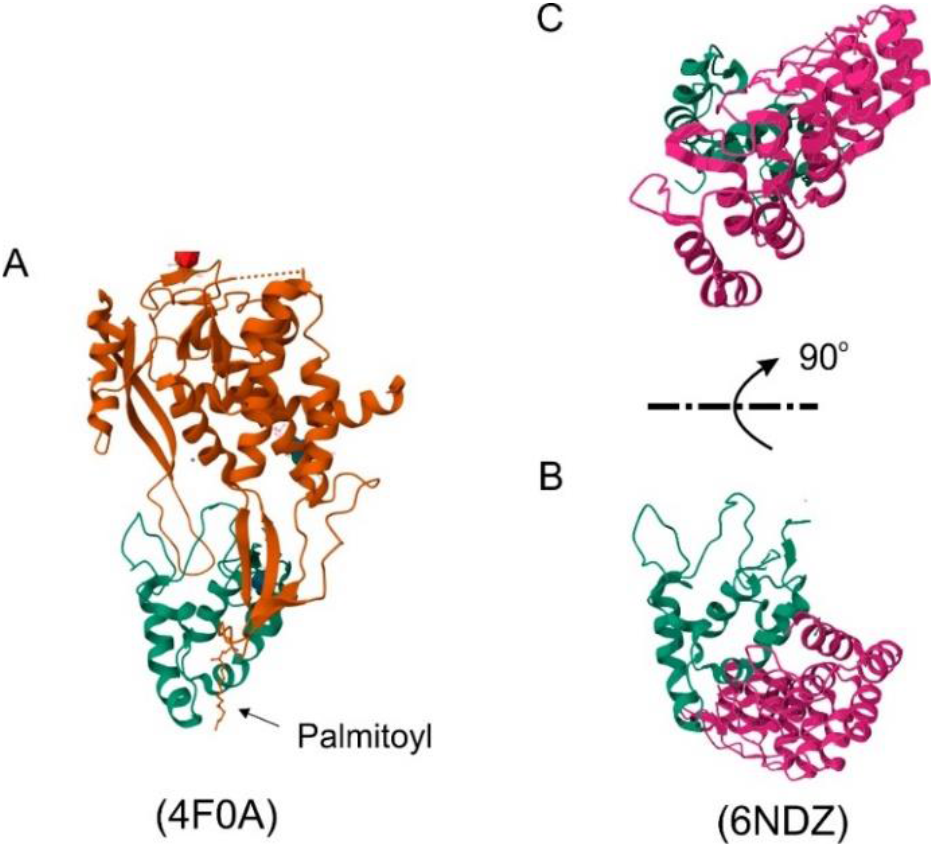
Comparison of Wnt-Fzd and NGS-Fzd complexes. (A) Structure of XWnt3a (orange) bound to the CRD of mFzd8 (green). Palmitoyl group on XWnt3a is highlighted by an arrow. (B) Structure of NGS (magenta) bound to mFzd8 (green), which is positioned in the same orientation as in A. (C) NGS-mFzd8 complex shown in B turned for 90 degree. PDB-IDs are shown in the brackets.

## SI Movies

**Movie S1.**
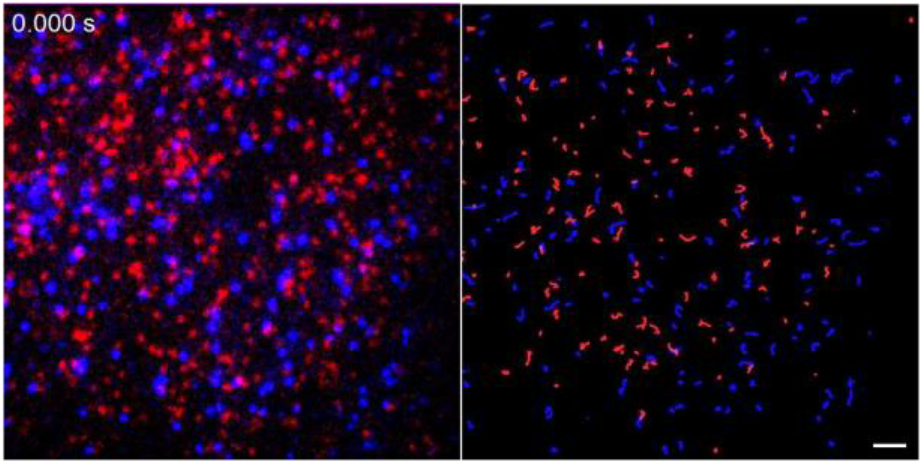
Two-color single-molecule tracking of Ror2 at the resting state. mXFPe-Ror2 was stably labeled by anti-GFP enhancer nanobodies conjugated with Dy647 (^Dy647^EN, blue) and Rho11 (^Rho11^EN, red). Left: Raw TIRF microscopy images recorded at video rate with 30 frames per second. Right: Single-molecule trajectories of ^Dy647^EN (blue) and ^Rho11^EN (red)-labeled mXFPe-Ror2. Scale bar: 5 µm.

**Movie S2.**
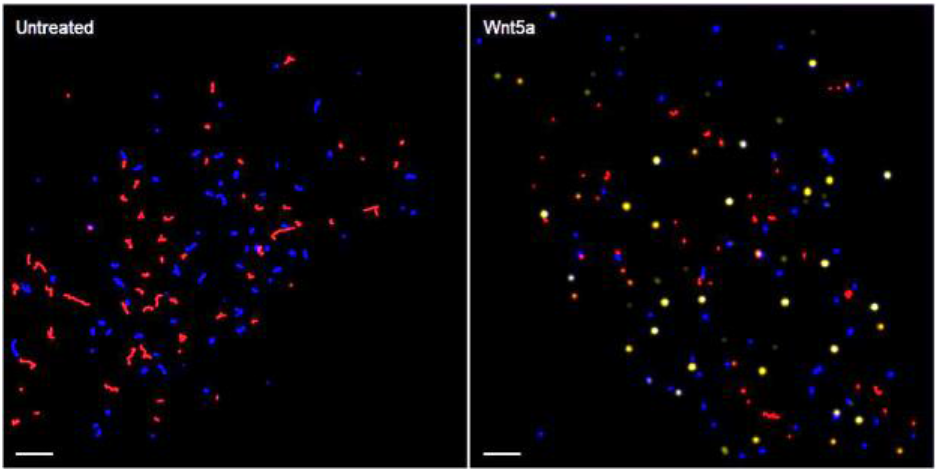
Two-color single-molecule tracking of Ror2 and Fzd8. mXFPe-Ror2 and mXFPm-Fzd8 were stably labeled with ^Rho11^EN and ^Dy647^MI nanobodies, respectively. Real-time playback at 30 frames per second. Scale bars: 5 µm. Left: Single molecule trajectories of Ror2 (red) and Fzd8 (blue) of an untreated cell. Right: Trajectories of Ror2 (red) and Fzd8 (blue) of another cell after Wnt5a treatment. Yellow dots mark the co-confined Ror2/Fzd8.

**Movie S3.**
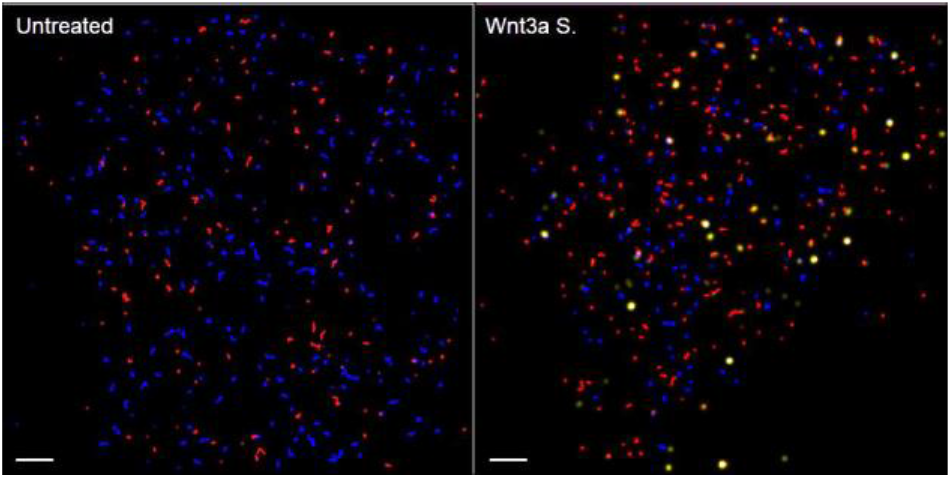
Two-*color* single-molecule tracking of Lrp6 and Fzd8. mXFPe-Lrp6 and mXFPm-Fzd8 were stably labeled with ^Rho11^EN and ^Dy647^MI nanobodies, respectively. Real-time playback at 32 frame-per-second. Scale bars: 5 µm. Left: Single molecule trajectories of Lrp6 (red) and Fzd8 (blue) of an untreated cell. Right: Trajectories of Lrp6 (red) and Fzd8 (blue) of another cell after Wnt3a surrogate treatment. Yellow dots mark the colocomoted Lrp6/Fzd8 within a distance of 150 nm.

**Movie S4.**
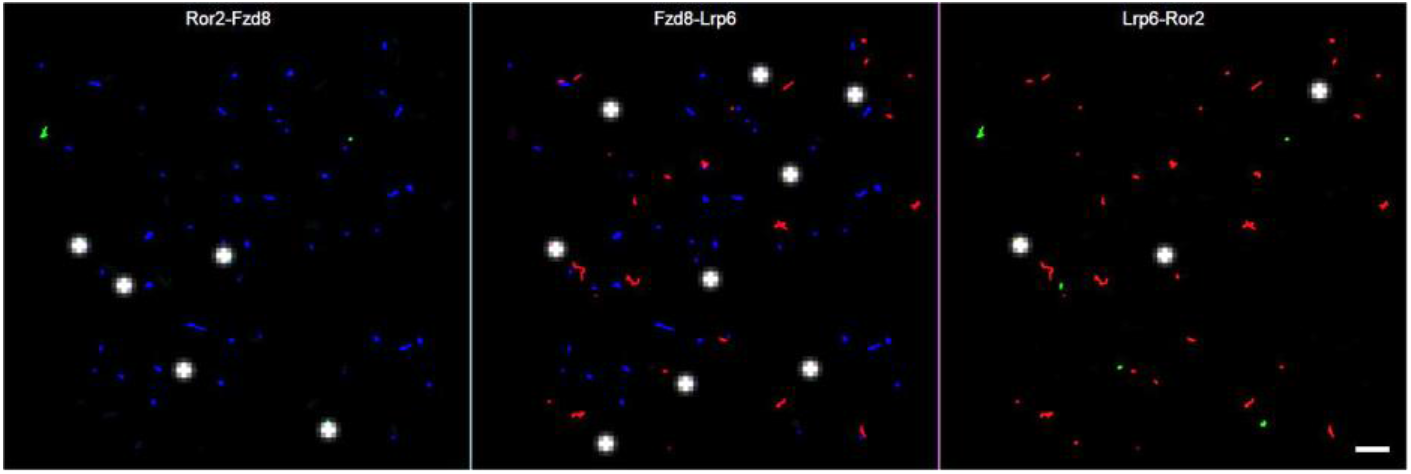
Three-*color* single-molecule tracking of Ror2, Lrp6 and Fzd8. Receptors were triple-color labeled as: mXFP-Ror2 by ^Dy752^EN (green), HaloTag-Lrp6 by ^TMR^HTL (red) and SNAP-Fzd8 by ^A647^SNAP Surface (blue). Wnt3a and Wnt5a were used for co-stimulation of canonical and noncanonical Wnt signaling. Real-time playback at 32 frame-per-second. Scale bar: 5 µm. Left: Single molecule trajectories of Ror2 (green) and Fzd8 (blue). Middle: Single molecule trajectories of Fzd8 (blue) and Lrp6 (red). Right: Single molecule trajectories of Lrp6 (red) and Ror2 (green). White dots mark co-confined co-receptors as indicated in the captions.

**Movie S5.**
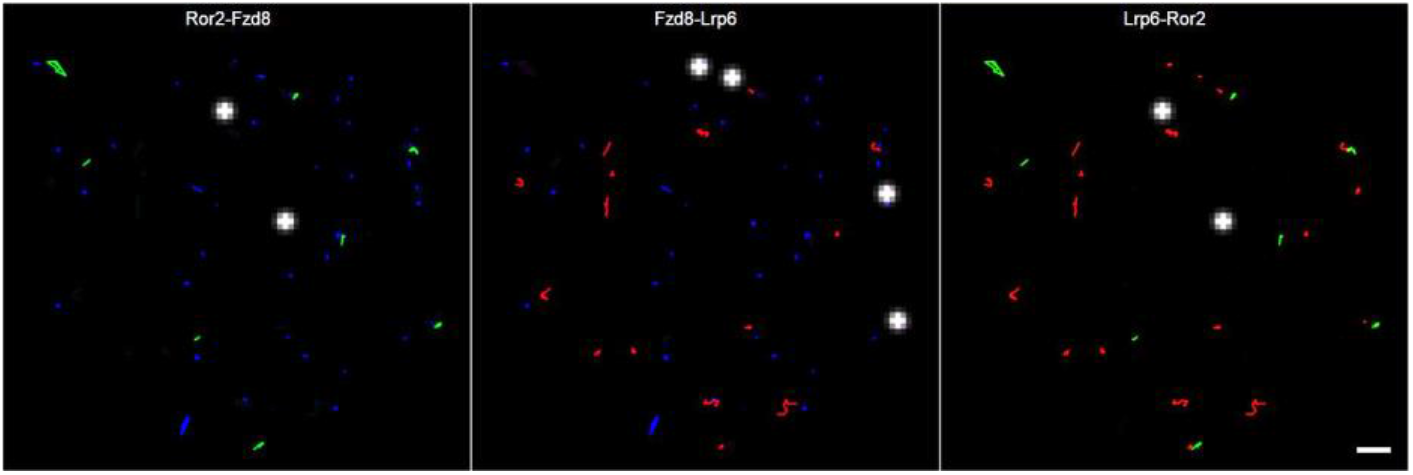
Three-*color* single-molecule tracking of Ror2, Lrp6 and Fzd8. Receptors were triple-color labeled as: mXFP-Ror2 by ^752^EN (green), Halo-Lrp6 by ^TMR^HTL (red) and SNAP-Fzd8 by ^A647^SNAP Surface (blue). Wnt3a surrogate and Wnt5a were used for co-stimulation of canonical and noncanonical Wnt signaling. Real-time playback at 32 frame-per-second. Scale bar: 5 µm. Left: Single molecule trajectories of Ror2 (green) and Fzd8 (blue). Middle: Single molecule trajectories of Fzd8 (blue) and Lrp6 (red). Right: Single molecule trajectories of Lrp6 (red) and Ror2 (green). White dots mark co-confined co-receptors as indicated in the captions.

**Movie S6.**
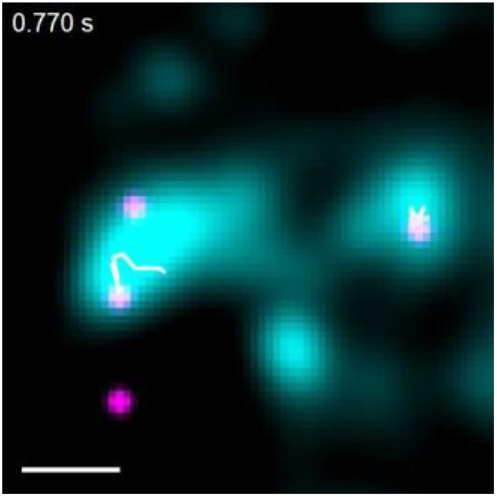
Confined diffusion of Lrp6 in a Wnt signalosome. HeLa cells stably expressing SNAP-Fzd8 (unlabeled) and HaloTag-Lrp6, which was labeled with ^Rho11^HD10 after stimulation with the Wnt surrogate NGS. The TALM image (cyan) outlining Wnt signalosomes was reconstructed from all localized Lrp6 detected within 11.7 min. Individual Lrp6 and their trajectories are shown in magenta. Scale bar: 200 nm.

**Movie S7.**
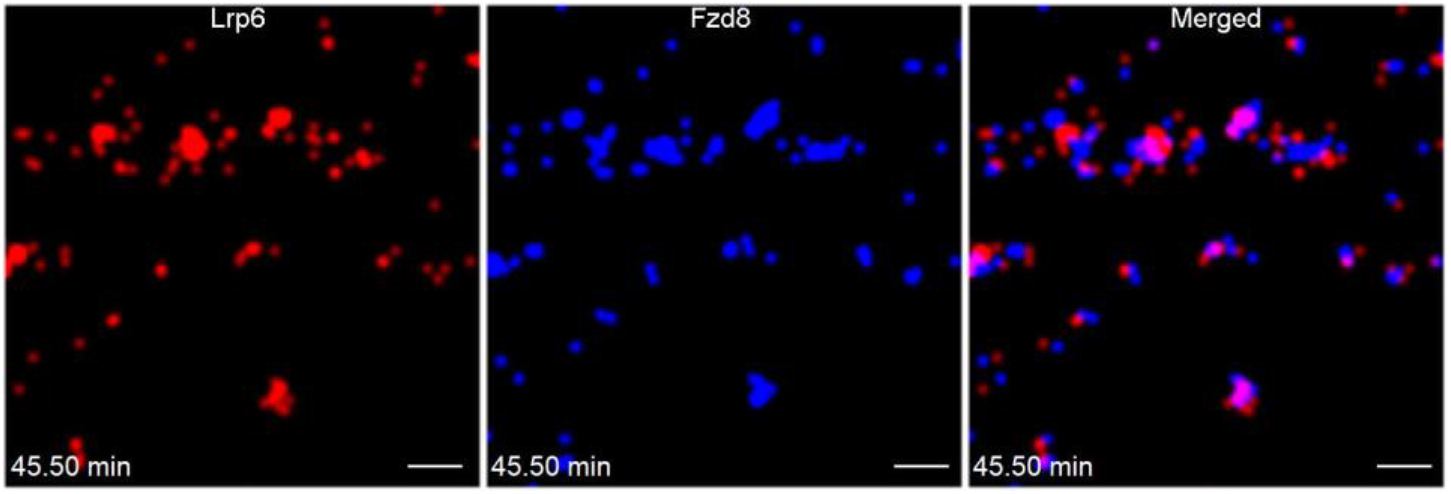
Time-lapse colocalization TALM images of co-localized Lrp6/Fzd8 in ROI1 of Fig. 4E. HaloTag-Lrp6 and CFP*HN*-Fzd8 were reversibly labeled with ^Rho11^HD10 and ^Dy647^EN. Agonist Wnt surrogate NGS was added for stimulation. The colocalized Lrp6 and Fzd8 was marked in red and blue, respectively, and their overlapping were shown in magenta in the ‘Merged’ channel. Time-lapse coloc-TALM images were reconstructed from colocalized Lrp6 and Fzd8 in the 2.9 min observation window (500 frames) and sliding with a 1.75 min step size (300 frames). Frame interval: 350 ms. Scale bars: 500 nm.

**Movie S8.**
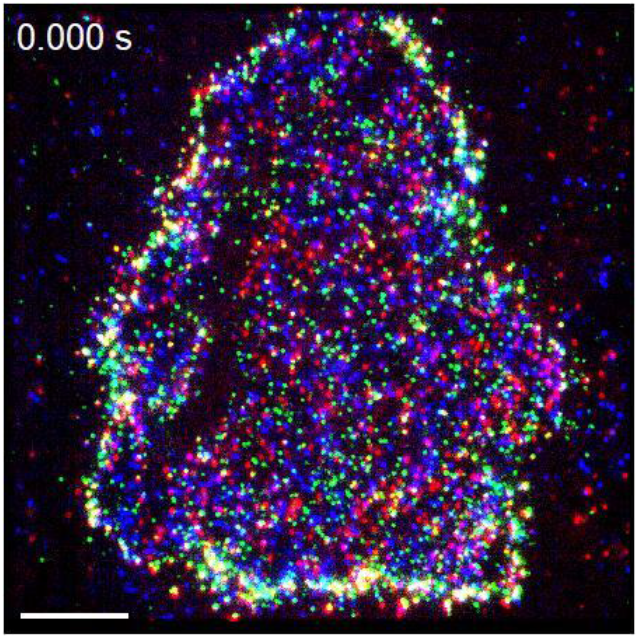
Three-color TIRFM images cell shown in FigS8A. ALFA-Ror2, Halo-Lrp6 and CFP*HN*-Fzd8 co-transfected in HeLa cell and reversibly labeled with ^Rho11^AnbAE, ^Dy647^HD10 and ^Dy752^EN, respectively. NGS and Wnt5a were added at 5.8 min and 47.1 min for activating canonical and noncanonical Wnt signaling. Images were taken at the end of joint stimulation (ca.100 min). Color coding: Lrp6 (red), Fzd8 (blue) and Ror2 (green). Scale bars: 10 µm.

**Movie S9.**
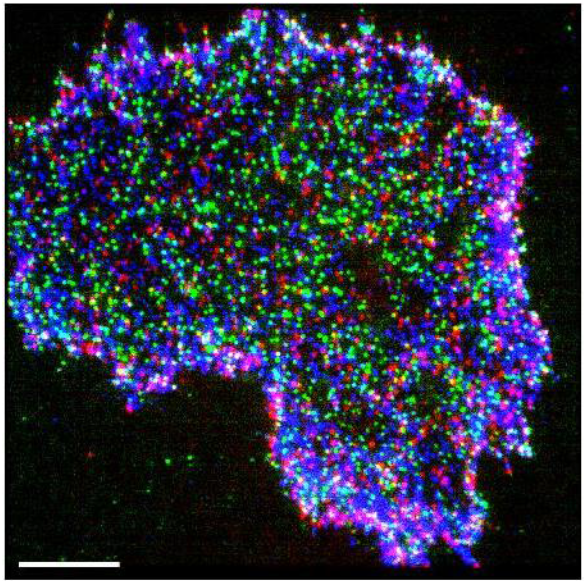
Three-color TIRFM images of cell shown in Fig S8E. ALFA-Ror2, Halo-Lrp6 and CFP*HN*-Fzd8 co-transfected in HeLa cell and reversibly labeled with ^Rho11^AnbAE, ^Dy647^HD10 and ^Dy752^EN, respectively. NGS and Wnt5a were added at 5.8 min and 47.1 min for activating canonical and noncanonical Wnt signaling. Images were taken at the end of joint stimulation (ca.100 min). Color coding: Lrp6 (red), Fzd8 (blue) and Ror2 (green). Scale bars: 10 µm. Real-time playback at 30 frame-per-second.

**Movie S10.**
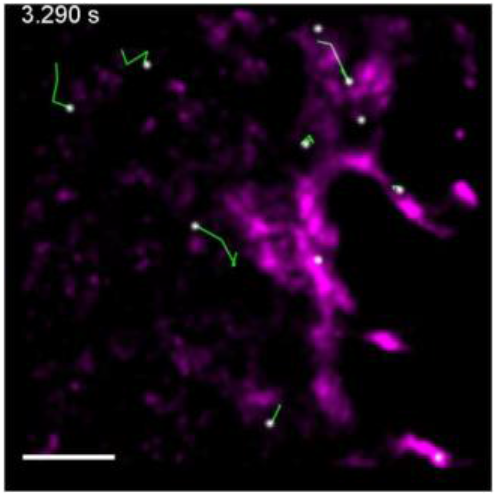
Transient confinement of Ror2 in the canonical Wnt signalosome. Trajectories of Ror2 merged with the colocalization TALM image of Lrp6/Fzd8 (magenta) after adding NGS but before Wnt5a (see Fig. 5G). ALFA-Ror2, Halo-Lrp6 and CFP*HN*-Fzd8 were reversibly labeled with ^Rho11^AnbAE, ^Dy647^HD10 and ^Dy752^EN. Ror2 is shown as white dot followed by green trajectories. Lrp6/Fzd8 co-clusters are shown in magenta in the background. Scale bar: 1 µm.

**Movie S11.**
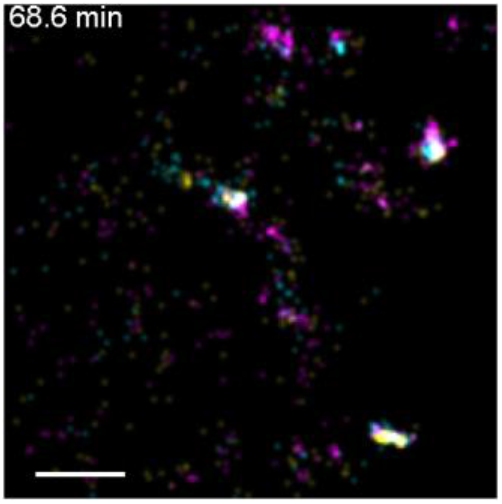
Time-lapse three-color coloc-TALM showing the formation of the common signalosome (see Fig. 5I). ALFA-Ror2, HaloTag-Lrp6 and CFP*HN*-Fzd8 were reversibly labeled with ^Rho11^AnbAE, ^Dy647^HD10 and ^Dy752^EN, respectively. NGS and Wnt5a were added sequentially for activating canonical and noncanonical Wnt signaling. Co-localized Lrp6/FZd8 (magenta), Fzd8/Ror2 (cyan) and Ror2/Lrp6 (orange) are shown. Time-lapse TALM images were obtained by accumulating colocalized receptors in a 300-frame observation window (1.4 min) and sliding at a stepwise of 300-frame (1.4 min). Frame interval: 280 ms. Scale bar: 1 µm.

**Movie S12.**
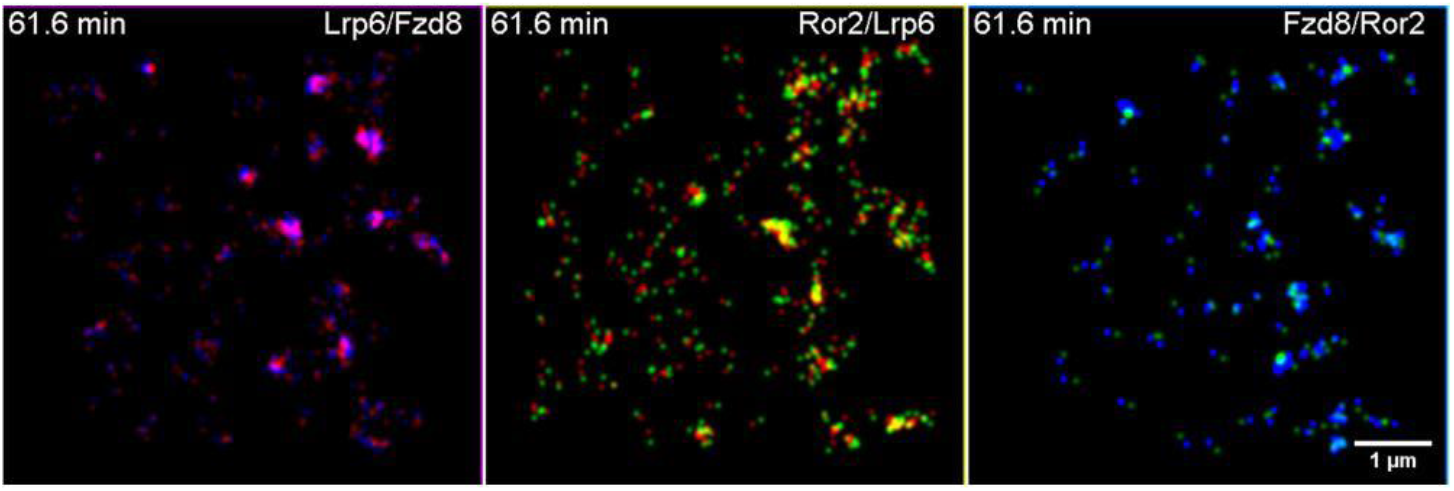
Three pairs time-lapse two-color coloc-TALM to deconvolve the formation of common signalosome (see Fig S10, i.e., ROI2 of Fig S8E). The time-lapse colocalization TALM images were reconstructed from colocalized Lrp6 (red)/Fzd8 (blue), Fzd8 (blue)/Ror2 (green) and Ror2 (green)/Lrp6 (red) in the 1.4 min observation window (300 frames) and sliding with a 1.4 min step (300 frames). Receptor pairs are indicated in each panel. Frame interval: 280 ms. Scale bar: 1 µm.

**Movie S13.**
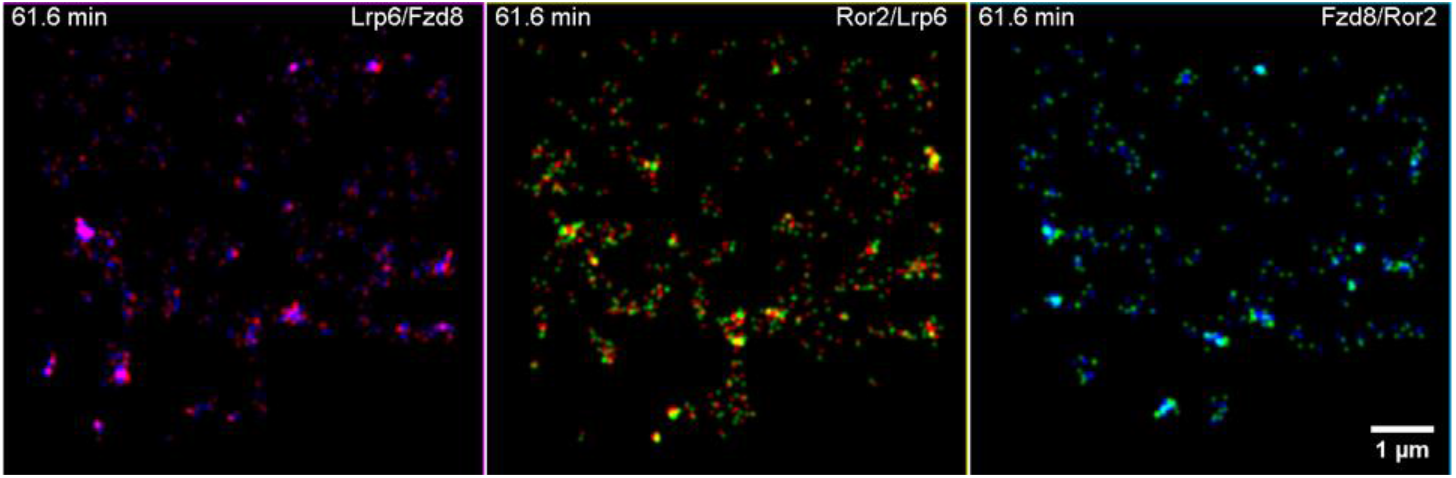
Three pairs time-lapse two-color coloc-TALM to deconvolve the formation of common signalosome (ROI1 of Fig S8E). Time-lapse colocalization TALM images were reconstructed by binary colocalizations of Lrp6 (red), Fzd8 (blue) and Ror2 (green) in the 1.4 min observation window (300 frames) and sliding with a 1.4 min step (300 frames). Frame interval: 280 ms. Receptor pairs are indicated in each panel. Scale bar: 1 µm.

## SI Tables

**Table S1:** Nanobodies, proteins and dyes.

| Protein (Denomination) / Dye | Source |
| --- | --- |
| HD10 anti-HaloTag nanobody (HD10) | This study |
| Anti-ALFA nanobody M63A/R65E (ALFAnbAE) | This study |
| Anti-GFP nanobody enhancer (EN) | Sotolongo Béllon et al., 2022 [42] |
| Anti-GFP nanobody minimizer (MI) | Sotolongo Béllon et al., 2022 [42] |
| Anti-maltose binding protein sybody (MSyb) | Zimmermann et al., 2018 [44] |
| mCFP-H164N (mCFPHN) | This study |
| ALFAtag-mEGFP | This study |
| Tagless mEGFP | Flesch et al., 2020[69] |
| Maltose binding protein (MBP) | Lisse, et al., 2011 [70] |
| HaloTag protein (Halo) | Lisse, et al., 2011 [70] |
| Wnt3a | R&D Systems |
| Wnt5a | R&D Systems |
| First generation Wnt surrogate (FGS) | Janda et al., 2017 [12] |
| Next generation Wnt surrogate (NGS) | Miao et al., 2020 [13] |
| Cy3B Maleimide | GE Healthcare |
| HTL-TMR | Promega |
| ATTO Rho11 Maleimide | ATTO-TEC |
| ATTO 643 Maleimide | ATTO-TEC |
| SNAP-Surface®647 | New England Biolabs* |
| DY-647P1 Maleimide | Dyomics GmbH |
| DY-752 Maleimide | Dyomics GmbH |

\* Discontinued and replaced by SNAP-Surface® 649, S9159S

**Table S2:**
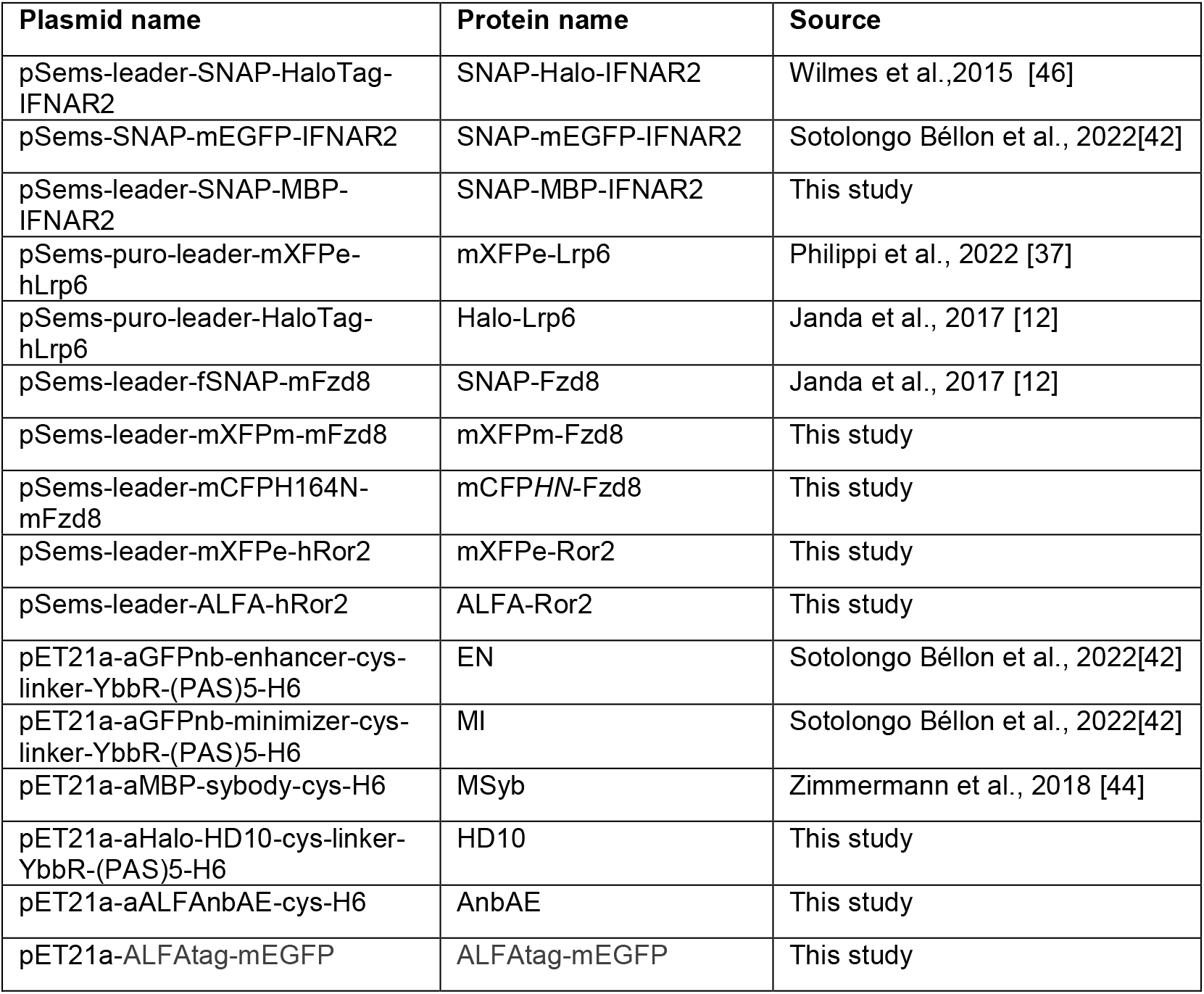
Recombinant DNA.

**Table S3:** Key materials, software and suppliers.

| Reagent and abbreviation | Source | Catalogue number |
| --- | --- | --- |
| *N*-2-Hydroxyethylpiperazine-*N*-2-ethane sulphonic acid (HEPES) | Carl Roth | 6763.3 |
| Ethylenediamine tetra-acetic acid (EDTA) | Carl Roth | 8040.2 |
| Dulbecco's Modified Eagle's Medium (MEM) | PAN-Biotech | P04-09500 |
| Dulbecco's phosphate buffered saline (PBS) | PAN-Biotech | P04-36500 |
| Fetal bovine serum (FBS) | PAN-Biotech | P30-3031 |
| Isopropyl-β-D-thiogalactopyranoside (IPTG) | Thermo Fisher Scientific | R0392 |
| DNase | Sigma-Aldrich/Merck | DN25 |
| Lysozyme | Sigma-Aldrich/Merck | L6876 |
| Urea | Carl Roth | 3941.2 |
| Protease inhibitor | Serva | 39106 |
| Chloramphenicol | Biomol | C4350.25 |
| Ampicilin | Biomol | 01503.25 |
| Trypsin | Capricorn Scientific | TRY-1B |
| ViaFect™ | Promega | E4982 |
| Accutase | Innovative Cell Technologies | AT104 |
| Linear polyethylenimine hydrochloride (PEI) | Polysciences | 24765-1 |
| 2-Amino-2-(hydroxymethyl)propane-1,3-diol (TRIS) | Sigma-Aldrich/Merck | 8.01464 |
| Glucose | Carl Roth | 3774.1 |
| Catalase | Sigma-Aldrich/Merck | C-40 |
| Glucose oxidase | Sigma Aldrich/Merck | 49180 |
| Methylviologen | Sigma Aldrich/Merck | 856177 |
| Ascorbic acid | Sigma Aldrich/Merck | 50817 |
| IWP2 | Sigma-Aldrich/Merck | I0536 |
| Poly-L-lysine poly(ethylene) glycol RGD (PLL-PEG-RGD) | You et al., 2014 [47] | N/A |
| CellSens Dimension | Olympus | 2.0 |
| CorelDraw | Corel | 2018 |
| Matlab R2013b | MathWorks | 2013 |
| BIAevaluation3.0 | BIACORE | 3.0 |
| OriginPro 9.0 | OriginLab | 9.0 |
| Software SLIMfast for single molecule analysis | Sotolongo Béllon et al., 2022[42] | N/A |
| Software TALM pair correlation analysis | You et al., 2014 [47] | N/A |
| ImageJ | Rasband et al., NIH | 1.54f (64bit) |

